# Sex and Region-Specific Disruption of Autophagy and Mitophagy in Alzheimer’s Disease: Linking Cellular Dysfunction to Cognitive Decline

**DOI:** 10.1101/2024.10.30.621097

**Authors:** Aida Adlimoghaddam, Fariba Fayazbakhsh, Mohsen Mohammadi, Zeinab Babaei, Amir Barzegar Behrooz, Farhad Tabasi, Teng Guan, Iman Beheshti, Mahmoud Aghaei, Daniel J Klionsky, Benedict C. Albensi, Saeid Ghavami

**Author notes:** Corresponding: Benedict C. Albensi, Nova Southeastern University, Ft. Lauderdale, FL, 33328, USA,; Saeid Ghavami, Department of Human Anatomy and Cell Science, Max Rady College of Medicine, Rady Faculty of Health Sciences, University of Manitoba, Winnipeg, MB, Canada. These authors of equivalent co-authorship (Fariba Fayazbakhsh, Mohsen Mohammadi). These authors have co-senior authorship (Benedict C. Albensi, Saeid Ghavami).

## Abstract

Macroautophagy and mitophagy are critical processes in Alzheimer’s disease (AD), yet their links to behavioral outcomes, particularly sex-specific differences, are not fully understood. This study investigates autophagy (LC3B-II, SQSTM1) and mitophagy (BNIP3L, BNIP3, BCL2L13) markers in the cortex and hippocampus of male and female 3xTg-AD mice, using western blotting, transmission electron microscopy (TEM), and behavioral tests (novel object recognition and novel object placement).

Significant sex-specific differences emerged: female 3xTg-AD mice exhibited autophagosome accumulation due to impaired degradation in the cortex, while males showed fewer autophagosomes, especially in the hippocampus, without significant degradation changes. TEM analyses demonstrated variations in mitochondrial and mitophagosome numbers correlated with memory outcomes. Females had enhanced mitophagy, with higher BNIP3L and BCL2L13 levels, whereas males showed elevated BNIP3 dimers. Cognitive deficits in females correlated with mitochondrial dysfunction in the cortex, while in males, higher LC3B-II levels associated positively with cognitive performance, suggesting protective autophagy effects.

Using machine learning, we predicted mitophagosome and mitochondrial numbers based on behavioral data, pioneering a predictive approach to cellular outcomes in AD. These findings underscore the importance of sex-specific regulation of autophagy and mitophagy in AD and support personalized therapeutic approaches targeting these pathways. Integrating machine learning emphasizes its potential to advance neurodegenerative research.

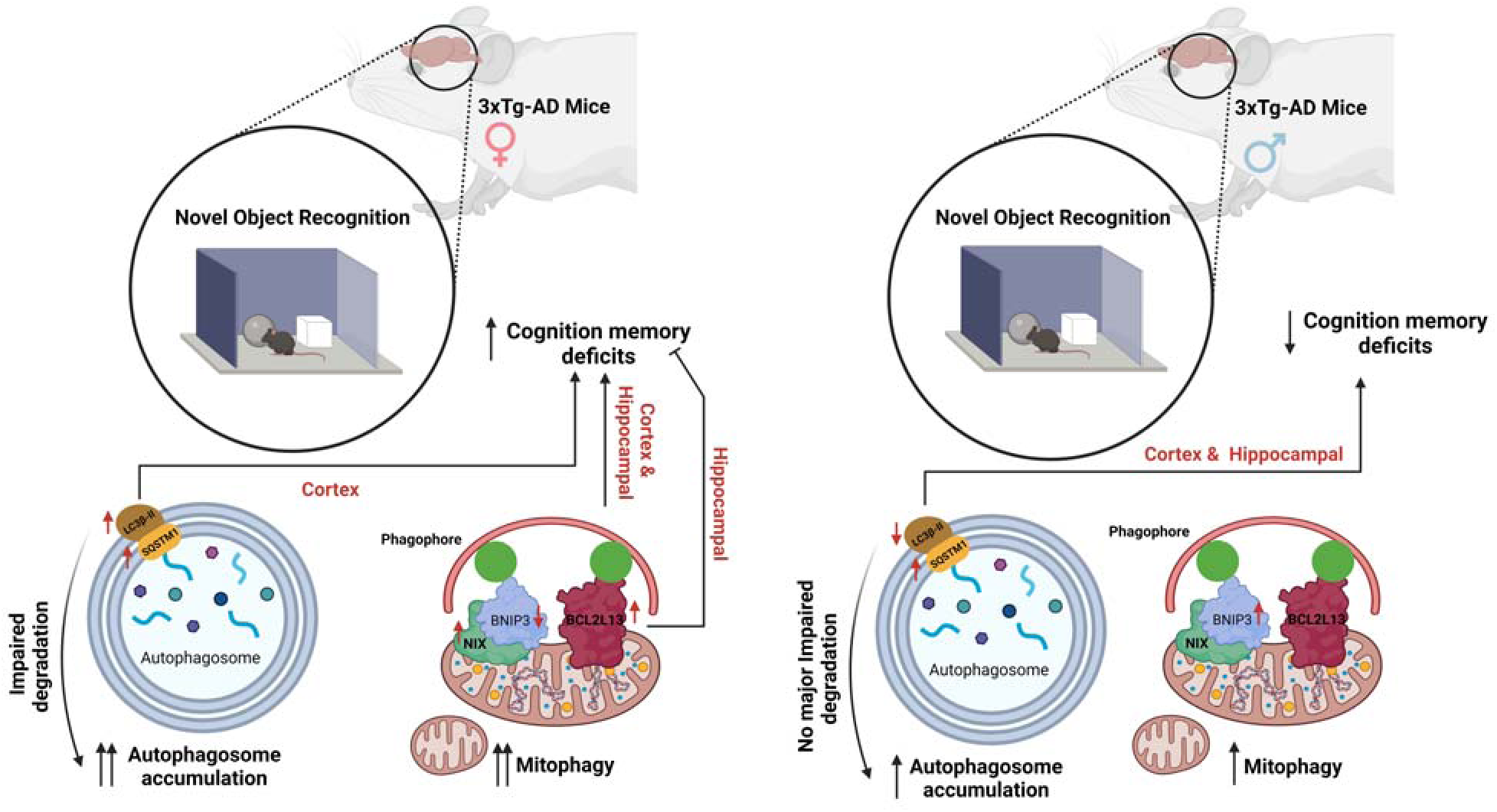

**Graphical Abstract:** Sex-specific differences in autophagy and mitophagy regulation in Alzheimer’s disease (AD) are highlighted. Female 3xTg-AD mice show autophagosome accumulation and cognitive deficits, while males exhibit variations in mitophagy markers and behavior.

## Introduction

Alzheimer’s disease (AD) is a devastating and irreversible neurodegenerative disorder that primarily affects older adults, leading to severe cognitive decline, memory deficits, and behavioral changes. Pathologically, AD is marked by the presence of amyloid-β (Aβ) plaques, hyperphosphorylated (p)-MAPT/tau tangles, and the accumulation of dysfunctional and/or damaged mitochondria in the brain (1–4). Collectively, in humans these pathological features result in synaptic and neuronal loss, particularly in the cerebral cortex and hippocampus, regions critical for cognitive function and memory (5, 6). Notably, cortical neurons are more vulnerable to mitochondrial disruption than hippocampal neurons in AD models (7).

A fascinating aspect of AD research is the significant influence of sex on disease risk and progression. Women exhibit a higher incidence and more severe pathology of AD, including impairments in working memory and spatial navigation tasks (8–10). Imaging studies further reveal a faster annual atrophy rate in female AD patients (11). Although the exact mechanisms behind these sex differences are not fully understood, factors such as cognitive reserve, genetics, and sex hormones are thought to contribute to differences in AD pathology and progression (12, 13).

Sex hormones, particularly estrogen, are pivotal in modulating autophagy and mitochondrial function (14, 15). Estrogen enhances basal autophagy levels, aiding in the clearance of MAPT/tau tangles and Aβ deposits (14). Moreover, female brains generally exhibit greater mitochondrial function compared to males (15). Consequently, the decline in estrogen levels in postmenopausal human females significantly reduces mitochondrial biogenesis and mitophagy in hippocampal neurons (16). This reduction in mitophagy is particularly concerning because hippocampal mitophagy is essential for sustaining neurogenesis and spatial memory (17).

A novel aspect of AD pathology is the impairment of mitophagy, a selective form of autophagy that removes damaged mitochondria through mitophagosomes, which then fuse with lysosomes to degrade the compromised mitochondria, maintaining cellular health and energy balance (18) (19). Mitophagy operates through various pathways, including those involving outer mitochondrial membrane proteins such as BNIP3 (BCL2/adenovirus E1B interacting protein 3), BNIP3L (BCL2/adenovirus E1B interacting protein 3-like), and BCL2L13 (BCL2 like 13) (20–22). These proteins interact with MAP1LC3/LC3-II via their LC3-interacting region/LIR, facilitating mitophagosome formation and expansion (22, 23).

BCL2-family proteins also play a dual role in regulating mitochondria-mediated cell death pathways (24). BNIP3, BNIP3L, and BCL2L13 show an interesting interplay between mitophagy and apoptosis, influencing various diseases, including AD (25–27). Notably, decreased levels of BNIP3L and BNIP3 in AD correlate with diminished mitophagy (28, 29).

We utilized novel object placement (NOP) and novel object recognition (NOR) behavioral tests to investigate the severe cognitive abnormalities associated with AD. These tests are powerful tools for assessing memory function, with NOP evaluating hippocampus-dependent spatial memory and NOR involving multiple brain regions, leveraging a mouse’s preference for novelty (30, 31). Exploring the relationship among these behavioral tests and also mechanisms of autophagy and mitophagy could provide groundbreaking insights into AD.

Recognizing that AD is characterized by autophagy and mitophagy impairment influenced by age, sex, and specific brain regions (16, 17, 32), we embarked on an innovative investigation to assess autophagy and mitophagy-related protein expression levels and also the number of mitochondria and mitophagosomes. Using the 3xTg-AD mouse model, widely recognized for its comprehensive representation of AD pathology (33), we aimed to correlate behavioral test performance in male and female 3xTg-AD mice with autophagy and mitophagy metrics in the cortex and hippocampus. This comprehensive analysis seeks to elucidate the role of mitophagy-related proteins and sex differences in AD pathogenesis, offering potential sex-specific therapeutic targets and paving the way for innovative personalized treatments.

## Materials and Methods

### Animal Models

Eleven-month-old male and female 3xTg and C57BL/6 mice (n=10 per genotype; n=5 female, n=5 male) were used. The 3xTg strain, carrying APPSwe, PSEN1M146V, and MAPT/TauP301L mutations, was maintained on a C57BL/6 background for over eight generations. Mice were housed in a pathogen-free facility at the St. Boniface Hospital Research Centre under a 12-h light/dark cycle with ad libitum access to food and water. All procedures were approved by the University of Manitoba Animal Care and Use Committee, following Canadian Council on Animal Care guidelines, and compliant with ARRIVE guidelines. A summary of the entire experimental procedure is presented in Scheme 1.

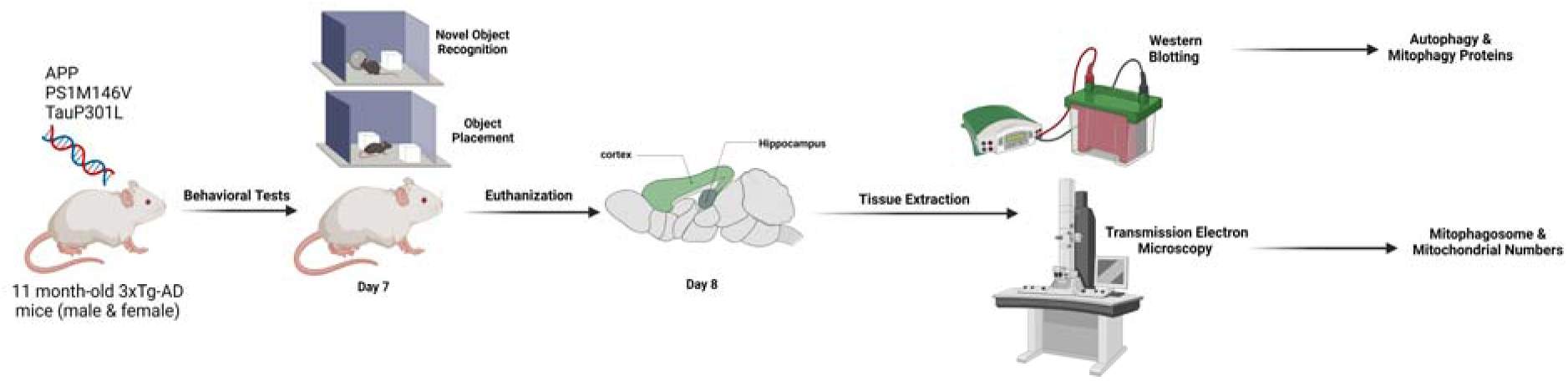

#### Summary of the method

In this study, 11-month-old male and female 3xTg-AD and C57BL/6 mice were subjected to behavioral tests, including Novel Object Recognition and Object Placement tests, to assess cognitive performance. Brain tissues were extracted for protein analysis using western blotting and for the quantification of mitochondria and mitophagosomes through transmission electron microscopy. These steps aim to examine autophagy and mitophagy in relation to AD progression. The entire experimental workflow is summarized in Scheme 1 for a comprehensive visual overview.

### Tissue Extraction

Mice were euthanized by decapitation under isoflurane anesthesia. Hippocampal and cerebral cortex (hereafter cortex) tissues were extracted, washed with PBS, and homogenized in ice-cold RIPA buffer containing phosphatase and protease inhibitors (Sigma-Aldrich). Homogenates were centrifuged, and supernatants were stored at -80°C. Protein concentrations were measured using a DC protein assay kit (Bio-Rad) as described previously (34).

### Western Blot

Hippocampal and cortical homogenates were prepared and subjected to SDS-PAGE using Criterion™ TGX Stain-Free™ gels (Bio-Rad). Proteins were transferred to nitrocellulose membranes and blocked in TBS-T with 5% BSA. The following primary antibodies were used: Total OXPHOS Rodent WB Antibody Cocktail (ab110413, Abcam), 1:1000 dilution; BNIP3L/NIX (D4R4B, Cell Signaling Technology), 1:1000 dilution; LC3B-II (048M4810V, Sigma-Aldrich), 1:4000 dilution; BCL2L13 (16612-1-AP, ProteinTech), 1:1000 dilution; BNIP3 (CS3769, Cell Signaling Technology), 1:1000 dilution; and SQSTM1/p62 (D1Q5S, Cell Signaling Technology), 1:1000 dilution. Secondary antibodies included goat anti-mouse IgG (H+L) and goat anti-rabbit IgG (H+L) (Jackson ImmunoResearch Laboratories), both at 1:2000 dilution. Enhanced chemiluminescence (ECL) was performed using the Clarity ECL kit (Bio-Rad) and imaged with the ChemiDoc™ MP system. Densitometry was quantified using ImageLab™ software, and normalized to total protein.

### Transmission Electron Microscopy (TEM)

Mice were perfused with PBS followed by 4% paraformaldehyde. Hippocampal and cortical tissues were fixed in Karnovsky’s fixative, post-fixed in osmium tetroxide, dehydrated in ethanol, and embedded in Epon resin. TEM imaging was performed using a Morgagni 268D electron microscope (Philips). Mitochondria and mitophagosomes were identified and quantified per cell. Mitochondria were identified by their double-membrane structure, typical size (0.5-1 μm), and intact cristae. Damaged mitochondria exhibited swollen or disrupted cristae. Mitophagosomes, larger than mitochondria, contained mitochondrial remnants within double-membrane vesicles, indicating degradation. Autophagosomes, also double-membraned, enclosed diverse cytoplasmic components marked for degradation. Multiple sections per sample were analyzed, and mitochondria, mitophagosomes, and autophagosomes were quantified per cell for statistical analysis.

### Novel Object Recognition and Novel Object Placement Tests

Behavioral testing was conducted on 11-month-old male and female 3xTg-AD and C57BL/6 mice (*n=10 per genotype; n=5 female, n=5 male*), seven days prior to euthanasia for tissue collection for western blotting and transmission electron microscopy (TEM). The Novel Object Recognition (NOR) and Novel Object Placement (NOP) **t**ests were designed to assess cognitive deficits related to memory and spatial recognition in Alzheimer’s disease (AD). The NOR test evaluates non-spatial forms of memory while the NOP test evaluates spatial memory. 1-**Familiarization (Habituation Phase):** In the initial phase, mice were exposed to an empty arena (*41 × 25 × 14 cm*) for 20 minutes, allowing them to become familiar with the testing environment without any objects present. This habituation established a baseline for exploratory behavior, ensuring that novelty alone, and not stress from the environment, would drive the results during testing. 2-**Training (Acquisition Phase):** During the **NOR** test, mice were introduced to two identical objects placed in the arena and allowed to explore them freely for 10 minutes. This phase familiarized the mice with the objects, allowing them to interact and establish a baseline for object recognition. 3-**Testing Phase (Novel Object Introduction):** After one hour, the **testing phase** was initiated by replacing one of the familiar objects with a **novel object**. Mice were allowed to explore the arena for 5 minutes, and their exploratory behavior was recorded. The **preference index** was calculated as the percentage of time spent exploring the novel object relative to the total exploration time. This process was repeated after a 24-hour interval, introducing a different novel object each time. 4-**Novel Object Placement (NOP) Test:** For the **NOP test**, the procedure was similar, with one key modification. After 5 minutes of training with two identical objects, one of the objects was moved to a new location in the arena. Mice were reintroduced to the arena for 5 minutes, and their interaction time with both the old and relocated objects was recorded. The **discrimination index** was calculated as (*time spent exploring the new place - time exploring the old place*)/total exploration time, assessing spatial memory and the mouse’s ability to detect changes in object placement (35).

### Machine Learning-Based Prediction

To explore the relationship between behavioral data and cellular outcomes, we applied machine learning techniques to predict mitophagosome and mitochondrial numbers using behavioral data as predictors. Data were collected from 20 mitophagosomes and 16 mitochondrial samples, with mitophagosome and mitochondrial counts as the target variables. Given the complexity of the data, we evaluated 11 different machine learning algorithms, including Linear Regression, Ridge Regression, Lasso Regression, ElasticNet Regression, Random Forest, Gradient Boosting, Support Vector Regression, Decision Tree, K-Neighbors Regression, Gaussian Process, and **XGBoost**. Since there is no standard way to determine the best algorithm for a specific dataset, we utilized a custom code to assess and compare the performance of all algorithms, with the best-performing algorithm selected for each experiment.

Leave-one-out cross-validation (LOOCV) was used for model validation, ensuring robust performance evaluation across the different algorithms. The final selected model provided the most accurate predictions based on R² values and Mean Absolute Error (MAE). For mitophagosome predictions, the best results were found in the hippocampus, with an R² of 0.57 and an MAE of 8.9, while in the **cortex**, the model yielded an R² of 0.27 and an MAE of 8.5. In predicting mitochondrial numbers, the highest accuracy was observed in the cortex (*R² = 0.47*, *MAE = 1.7*), whereas predictions for the hippocampus were less accurate (*R² = 0.001*, *MAE = 2.56*).

It’s important to note that only two behavioral features were used in this analysis, limiting the insights provided by the feature coefficients. Moreover, due to the distinct mechanisms employed by each algorithm for determining feature importance, a direct comparison of feature coefficients across models was not feasible. The selection of the best-performing algorithm was solely based on accuracy comparisons, and each algorithm was chosen for its specific predictive capability in different regions of the brain.

### Statistical Analysis

All statistical analyses were performed using SPSS software (version 17.0, SPSS Inc., Chicago, IL, USA), and graphs were drawn using GraphPad Prism 8.0.1 software (GraphPad Software, La Jolla, CA, USA). Statistical analysis of behavioral performance was conducted using a two-way analysis of variance (ANOVA). Spearman’s correlation test was applied to correlate the behavioral findings obtained in the NOP and NOR tests with mitochondria and mitophagosome numbers and BCL2-family protein expression in the cortex and hippocampus of female and male 3xTg-AD mice. All P-values are presented as two-tailed, and P < 0.05 was considered statistically significant.

## Results

### Sex-Specific Autophagosome Accumulation and Degradation in Alzheimer’s Disease

Autophagy plays a critical role in neurodegenerative diseases such as Alzheimer’s by regulating the removal of damaged proteins and organelles (36). We assessed key autophagy markers in 3xTg-AD mice to investigate potential sex-specific differences in AD autophagy pathways. We measured the amount of the phosphatidylethanolamine-conjugated form of MAP1LC3B/LC3B, **LC3B-II (**Figure 1A), an indicator of autophagosomes. In female 3xTg-AD mice, **LC3B-II** levels in the cortex were significantly elevated compared to controls (P < 0.05; Figure 1B), indicating an accumulation of autophagosomes, which could reflect autophagy induction or a block in degradation/autophagosome turnover. In male 3xTg-AD mice, **LC3B-II** levels were reduced in both the cortex and hippocampus as compared to controls, with a significant decrease in the hippocampus (P < 0.05; Figure 1C), suggesting fewer autophagosomes. The autophagic receptor and substrate **SQSTM1**, which marks autophagosome degradation, is depicted in Figure 1D through western blot analysis. Higher **SQSTM1** levels in the cortex (P < 0.05; Figure 1E) suggest impaired autophagosome degradation as the cause of LC3B-II accumulation. **SQSTM1** levels in males did not differ significantly from controls (P > 0.05; Figure 1F), indicating no major impairment in autophagosome degradation. These findings suggest sex-specific alterations in autophagy regulation in AD, with females showing autophagosome accumulation in the cortex due to impaired degradation, while males exhibit fewer autophagosomes, particularly in the hippocampus, without significant changes in degradation.

**Figure 1.**
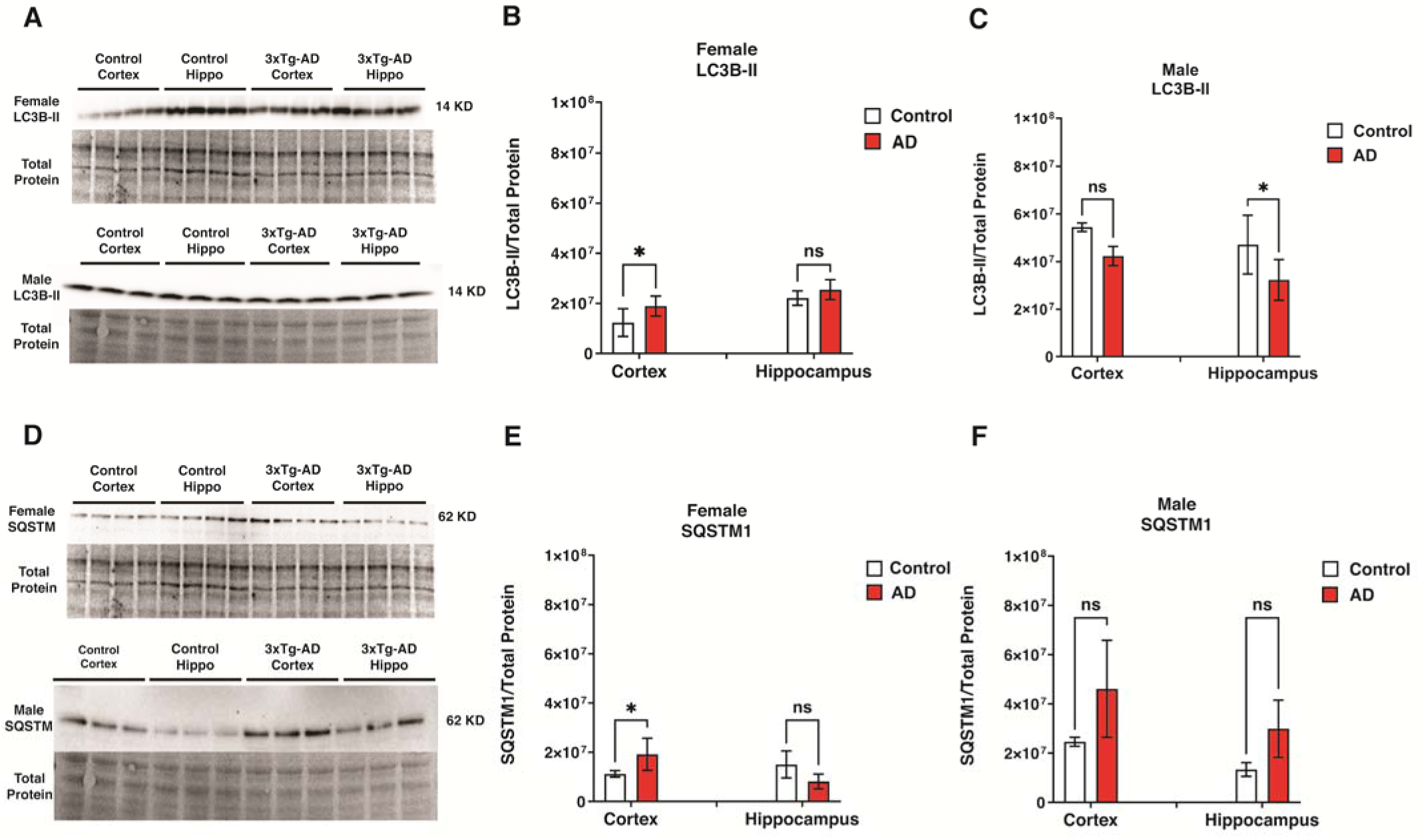
Autophagosome Status in Control and 3xTg-AD Mice. **(A, D)** Representative western blot bands showing LC3 and SQSTM1 protein levels in male and female mice. **(B, C)** Quantification of LC3B-II protein levels in the cortex and hippocampus of female and male mice. **(E, F)** Quantification of SQSTM1 protein levels in female and male mice. Results show significantly lower autophagosome degradation (SQSTM1) and higher autophagosome marker (LC3B-II) in the cortex of 3xTg-AD females (P < 0.05). In 3xTg-AD males, there was a significant reduction in the autophagosome marker (LC3B-II) in the hippocampus (P < 0.05). In contrast, no significant differences were found in autophagosome degradation (SQSTM1) in both the cortex and hippocampus (P > 0.05). Statistical analyses were performed using a two-way ANOVA test. *P < 0.05; **P < 0.01. Data are presented as mean ± SD, with n = 5 per group (n = 5 females and n = 5 males).

### Sex-Specific Alterations in Mitophagy-Related BCL2 Protein Expression in Alzheimer’s Disease

The BCL2 family proteins BNIP3L, BNIP3, and BCL2L13 play a key role in regulating mitophagy (37–39), a process crucial for maintaining mitochondrial integrity. In this study, we explored sex-specific differences in the expression of these proteins in Alzheimer’s disease to understand their potential impact on mitophagy.

To this end, we measured cortical and hippocampal levels of outer mitochondrial membrane proteins **BNIP3L**, **BNIP3**, and **BCL2L13** using western blot analysis to examine potential changes in mitophagy activity (Figure 2). **BNIP3L** dimers more efficiently recruit phagophores than the monomeric form (40). **The BNIP3L** monomer levels were significantly higher in the cortex (P < 0.0001) and hippocampus of 3xT-AD female mice (P < 0.001; Figure 2C) but not in males (Figure 2D). Female 3xTg-AD mice also showed lower **BNIP3L** dimer levels in the hippocampus (P < 0.001) and higher levels in the cortex (P < 0.001; Figure 2E) compared to controls. In the male group, no significant differences in **BNIP3L** dimer levels were found between control and 3xTg-AD (Figure 2F).

**Figure 2.**
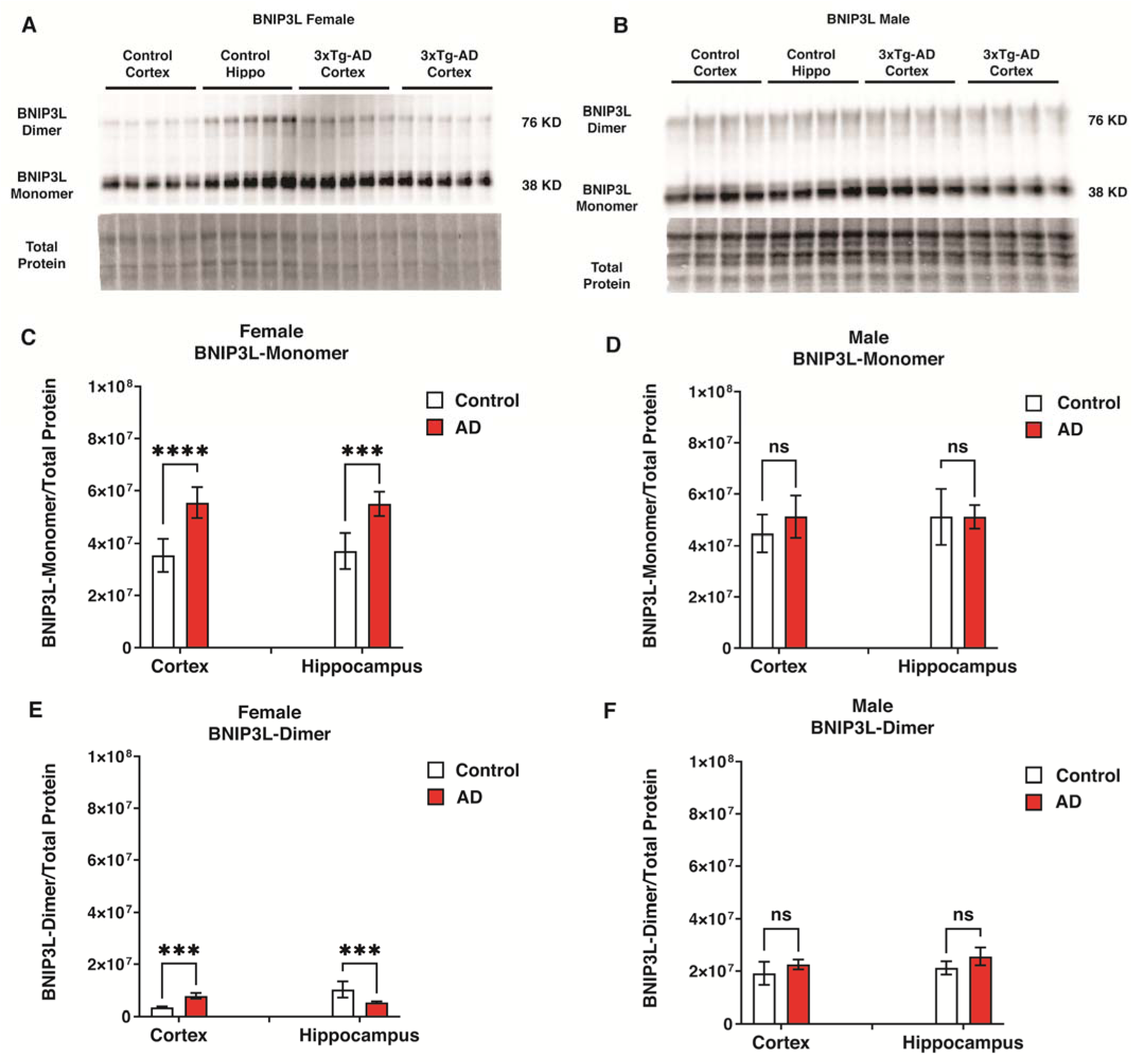
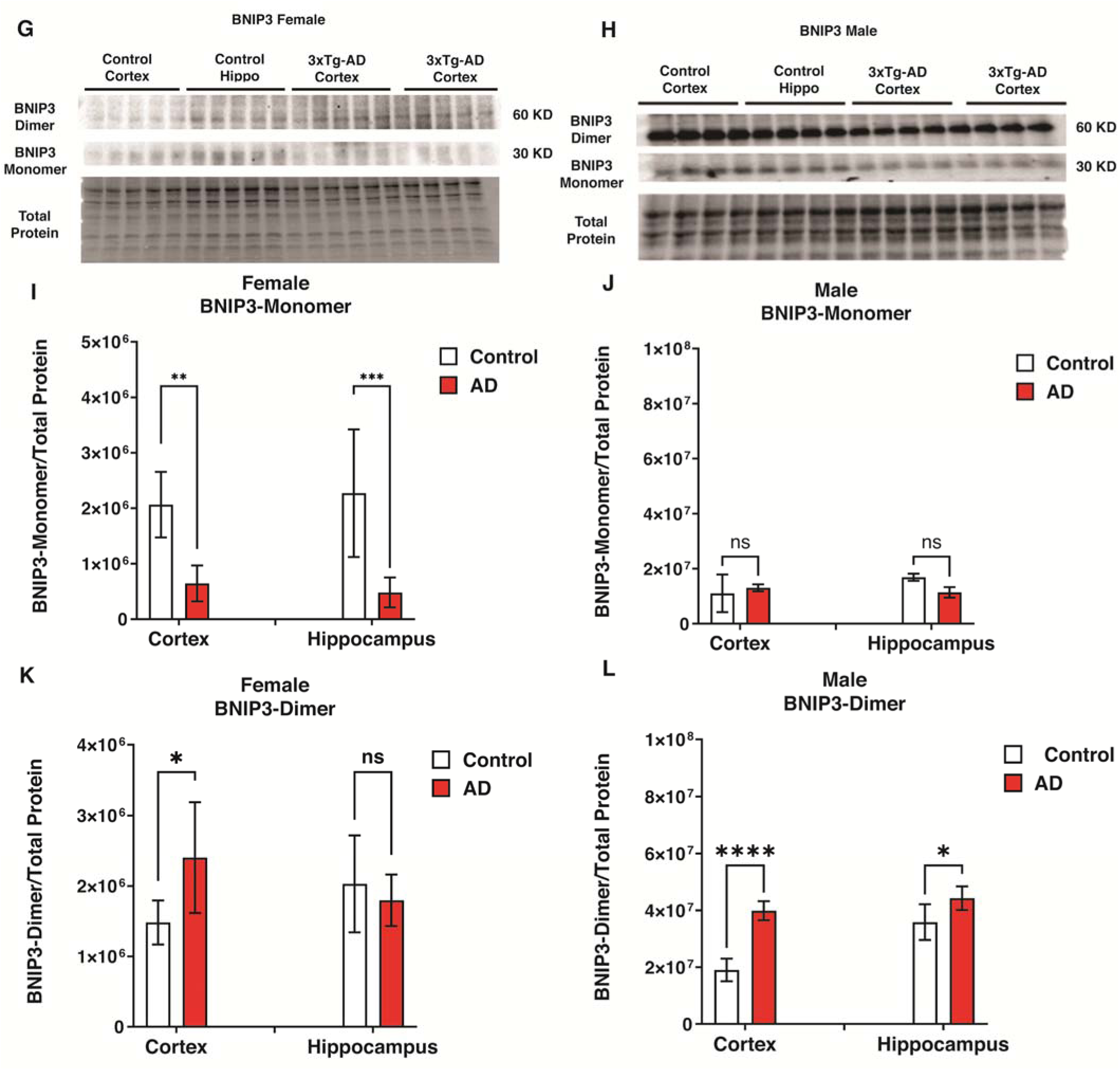
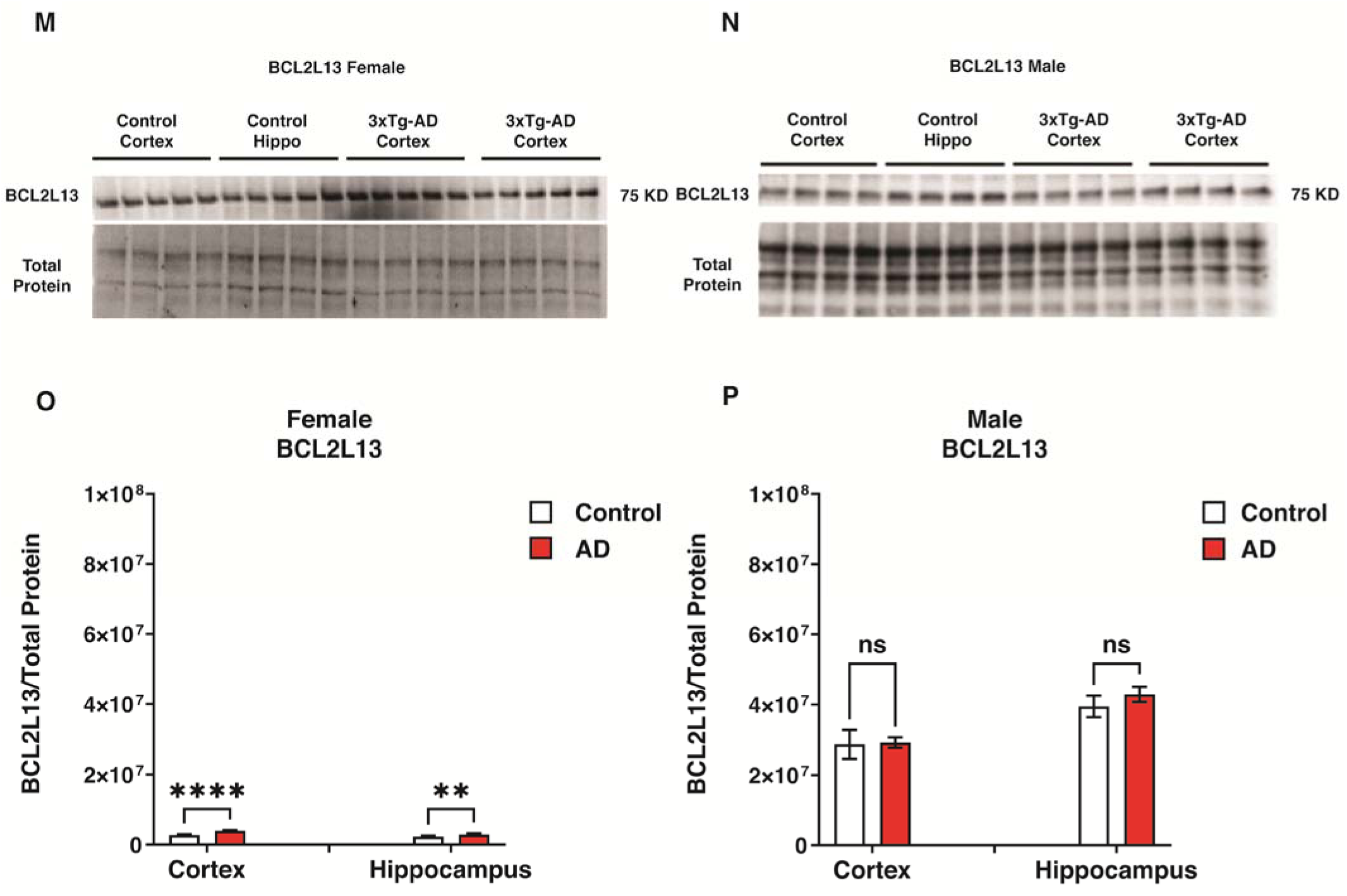
Mitophagy-related BCL2-Family Protein Status in Control and 3xTg-AD Mice. **(A, D)** western blot analysis of BNIP3L monomer and dimer in the cortex and hippocampus of male and female 3xTg-AD and control mice. **(B, E)** Quantification of BNIP3L monomer expression levels in females and males, respectively. In female 3xTg-AD mice, a significant increase in BNIP3L monomer expression was observed in the cortex (P < 0.0001) and hippocampus (P < 0.001). **(C, F)** Quantification of BNIP3L dimer levels in female and male mice, respectively. Comparing the control and 3xTg-AD groups, a significantly higher BNIP3L dimer level was found in the cortex (P < 0.01), while a lower level was observed in the hippocampus of 3xTg-AD females (P < 0.01). No significant differences in BNIP3L monomer or dimer levels were found between control and AD males. **(G, J)** Western blot analysis of BNIP3 monomer and dimer in female and male mice. **(H, K)** Quantification of BNIP3 monomer expression levels in females and male mice, respectively. BNIP3 monomer expression was significantly lower in the cortex (P < 0.01) and hippocampus (P < 0.001) of 3xTg-AD females than in control, while no significant differences were found in male groups. **(I, L)** Quantification of BNIP3 dimer levels in females and males, respectively. Our investigation showed higher BNIP3 dimer level in the cortex (P < 0.0001) and the hippocampus (P < 0.05) of the 3xTg-AD male group and also in the cortex (P < 0.05) of 3xTg-AD female compared to the control. **(M, O)** Western blot analysis of BCL2L13 in female and male mice. **(N)** Quantification of the expression levels of BCL2L13 in the cortex and hippocampus of female 3xTg-AD and control mice. BCL2L13 expression in the cortex (P < 0.0001) and the hippocampus (P < 0.01) of 3xTg-AD females is higher than in control. **(P)** Quantification of the expression levels of BCL2L13 in the cortex and hippocampus of male 3xTg-AD and control mice. No significant differences were found in male groups. Statistical analyses were performed using a two-way ANOVA test. *P < 0.05; **P < 0.01; ***P < 0.001; ****P < 0.0001. The results are presented as the mean ± SD, n = 5 per group (n = 5 females and n = 5 males).

Similar to **BNIP3L**, **BNIP3** also dimerizes, which is important for functions including lysosomal delivery(40, 41). Figures 2G and 2H show western blot analysis of BNIP3 monomer and dimer in female and male mice. The expression of the **BNIP3** monomer was significantly lower in the cortex (P < 0.01) and hippocampus (P < 0.001) of the 3xTg-AD female group compared to the control group (Figure 2I). In contrast, no significant differences were observed in the male groups (Figure 2J). In our study of the 3xTg-AD female group, we found higher levels of the BNIP3 dimer in the cortex (P < 0.05; Figure 2K) compared to the control group. However, we did notice higher levels of the BNIP3 dimer in the cortex (P < 0.0001) and hippocampus of male 3xTg-AD mice (P < 0.05; Figure 2L).

The western blot analysis of BCL2L13 in female and male mice is illustrated in Figures 2M and 2N. In female mice, **BCL2L13** expression in the cortex (P < 0.0001) and the hippocampus (P < 0.01) was higher than in the control (Figure 2O). No significant differences were found in the male groups (Figure 2P).

These findings reveal significant sex-specific differences in the expression of BCL2-family proteins that regulate mitophagy, suggesting potential changes in mitophagy activity. In females, elevated BNIP3L and BCL2L13 levels in the cortex and hippocampus point to enhanced mitophagy, while males exhibited increased BNIP3 dimer levels, particularly in the cortex. These results indicate that sex may influence mitochondrial maintenance and degradation pathways in Alzheimer’s disease.

### Sex-Specific Differences in Mitochondrial and Mitophagosomal Dynamics in Alzheimer’s Disease

Maintaining mitochondrial homeostasis is crucial in neurodegenerative diseases such as AD (42). In this study, we aimed to examine sex-specific differences in mitochondrial content and mitophagy activity by quantifying mitochondria and mitophagosome numbers in 3xTg-AD mice using TEM.

Using TEM, we identified the number of mitophagosomes and mitochondria in the cortex and hippocampus of control and 3xTg-AD groups. The number of mitochondria in the cortex of male 3xTg-AD mice was significantly lower than in the control group (P < 0.01; Figure 3B). Additionally, we observed a significantly higher number of mitochondria in the cortex (P < 0.001) and hippocampus (P < 0.0001) of female controls compared to the 3xTg-AD group (Figure 3C). Our assessment indicated the number of mitophagosomes was significantly higher in the cortex (P < 0.0001) and hippocampus (P < 0.0001) of male 3xTg-AD mice compared to the control group (Figure 3D). Similarly, a significant increase in the number of mitophagosomes in the cortex of female 3xTg-AD mice compared to the control group (P < 0.01; Figure 3E). Our findings exhibited similar trends in both the cortex and hippocampus. However, there were significant differences between the two sexes. For example, the number of mitochondria in male mice showed no statistical difference in the hippocampus and a relatively small difference in the cortex. In contrast, female mice displayed much larger differences in both locations. Conversely, the number of mitophagosomes in the hippocampus was substantially higher in the 3xTg-AD male mice compared to the control, whereas there was no statistical difference in the female mice. These results suggest that mitophagy may be differentially regulated between males and females in AD.

**Figure 3.**
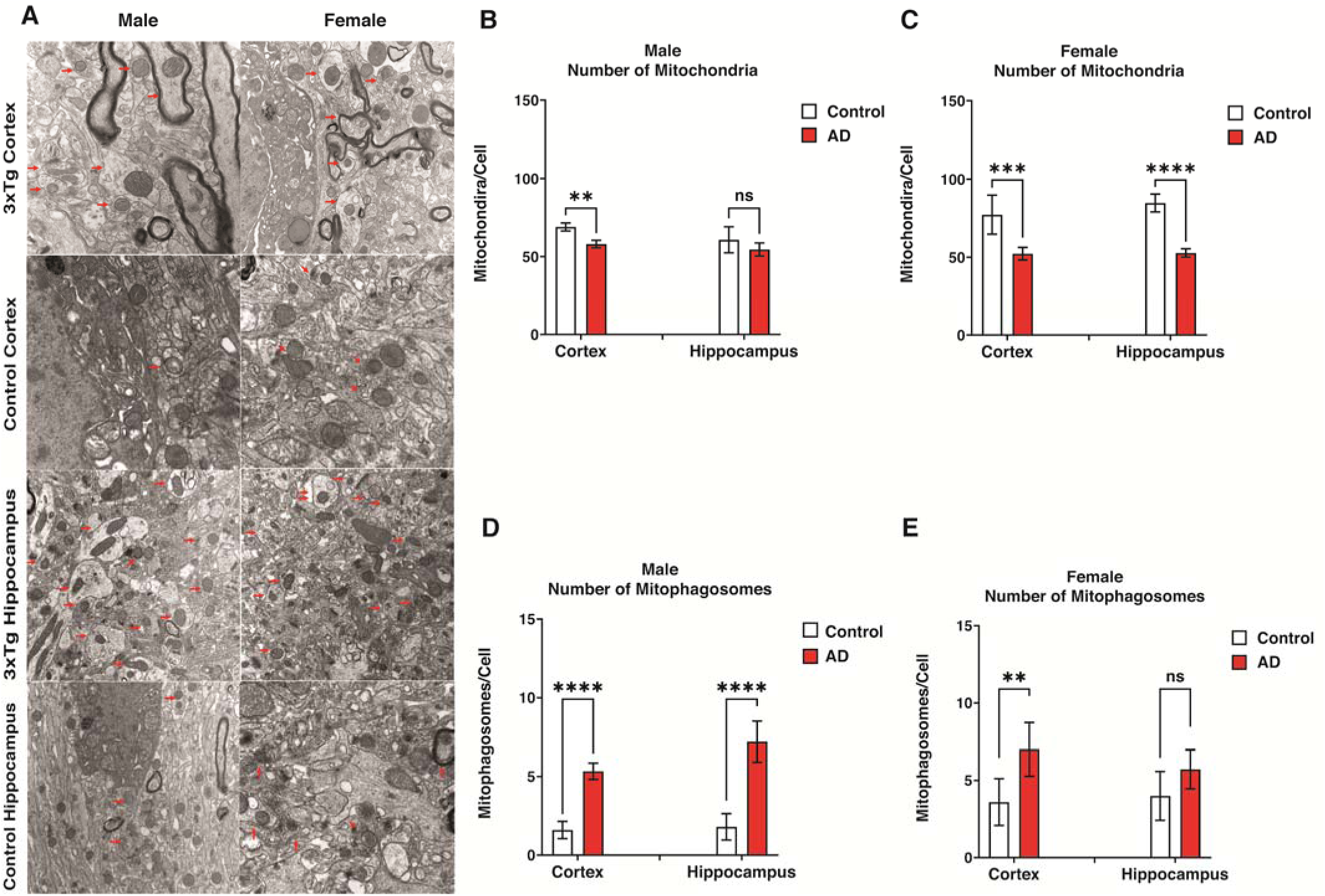
Transmission Electron Microscopy Reveals Mitochondrial and Mitophagosome Changes in 3xTg-AD Mice in the Cortex and Hippocampus. **(A)** Transmission electron microscopy images of the cortex and hippocampus in male and female control and 3xTg-AD mice. Arrows indicate mitophagosomes and mitochondria. **(B, C)** Quantification of mitochondria in the cortex of female and male mice, respectively. The lower number of mitochondria in the cortex (P < 0.001) and hippocampus (P < 0.0001) of 3xTg-AD females indicates potentially higher mitophagy rates in the AD compared to the control female group. In males, the potential mitophagy rate in the cortex of the 3xTg-AD group is significantly higher than the control (P < 0.01). **(D, E)** Quantification of mitophagosomes in the cortex of female and male mice, respectively. A significant increase in the mitophagosome number of the cortex in male 3xTg-AD compared to the control (P < 0.01) indicates a potentially higher mitophagy rate (or lower mitophagosome degradation) in male 3xTg-AD compared to the control. Similarly, in females, the number of mitophagosomes is significantly increased in the cortex (P < 0.0001) and the hippocampus (P < 0.0001) of 3xTg-AD compared to the control group, indicating a potentially higher mitophagy rate (or lower mitophagosome degradation) in the cortex and hippocampus of AD compared to control. Statistical analyses were performed using a two-way ANOVA test. *P < 0.05; **P < 0.01; ***P < 0.001; ns, non-significant. The results are presented as the means ± SD.

### Sex-Specific Cognitive Deficits and Correlations with Autophagy and Mitophagy proteins in 3xTg-AD Mice

Progressive cognitive decline is the typical clinical feature of AD (1). Mitochondrial dysfunction and mitophagy failure are possible mechanisms of cognitive deficits in AD (4). Therefore, we next examined the contribution of neuronal mitophagy on learning and memory impairments in 3xTg-AD mice. Spatial and non-spatial memory was investigated in 3xTg-AD mice compared to control mice using novel object placement (NOP for spatial memory) and novel object recognition (NOR for non-spatial memory) behavioral tests. During the NOR test, female 3xTg-AD mice exhibited less preference for the novel object compared to control females (P < 0.0001) and also 3xTg-AD males (P < 0.01), but no significant difference was observed between male 3xTg-AD mice and the control group (P > 0.05) (Figure 4A). In the NOP test, there were no significant differences in preference index between female or male 3xTg-AD mice and controls (P > 0.05) (Figure 4B). These results indicate non-spatial memory deficits in female 3xTg-AD mice, but not in male, as demonstrated by the NOR test.

**Figure 4.**
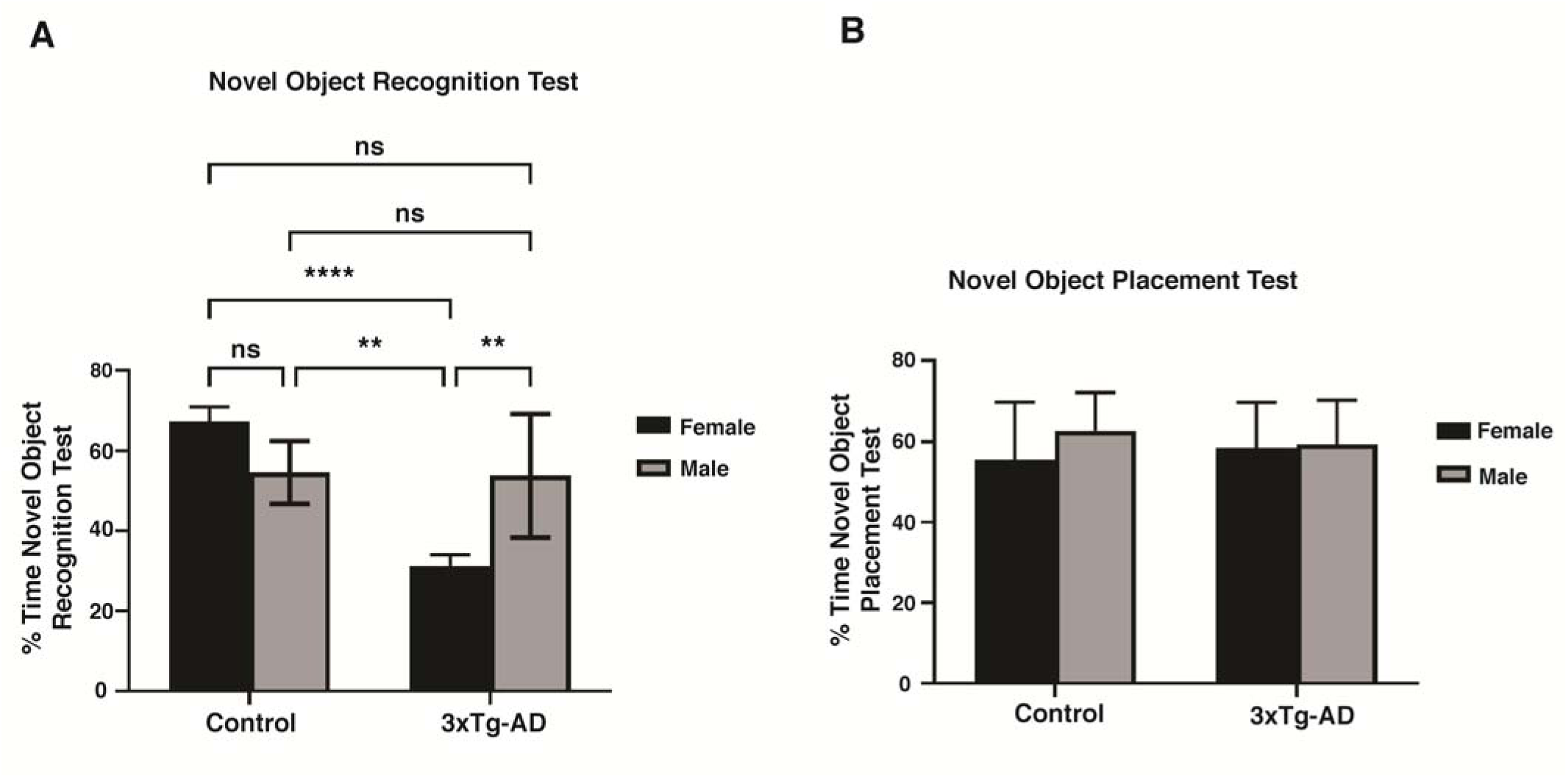
Behavioral Assessment of Cognitive Performance in 3xTg-AD Mice Using Novel Object Recognition (NOR) and Novel Object Placement (NOP) Tests. **(A)** Memory impairment (NOR test) comparison between female and male control and 3xTg-AD mice. Females exhibited significant non-spatial memory deficits compared to 3xTg-AD males (P < 0.01) and the female control group (P < 0.0001). **(B)** Spatial memory impairment (NOP test) comparison between female and male control and 3xTg-AD mice. Our analysis showed no significant differences in spatial memory between male and female 3xTg-AD mice and controls. Statistical analyses were performed using a two-way ANOVA test. **P < 0.01; ****P < 0.0001; ns, non-significant. Data are presented as mean ± SD, n = 5 per group (n = 5 females, n = 5 males).

We further evaluated the correlations of behavior findings with expression levels of the autophagy-related proteins LC3B-II and SQSTM1 in the cortex and hippocampus of 3xTg-AD mice. The results of Spearman’s correlation test showed that the autophagosome (LC3B-II) levels in the cortex of male 3xTg-AD mice were negatively correlated with the NOP test (r = - 1.000, P = 0.010; Figure 5A), but positively correlated with the preference index of the novel object in the NOR test (r = 1.000, P = 0.010; Figure 5B). In contrast, in females, no associations were found between autophagosome levels and NOR (Figure S1A) or NOP tests (Figure S1B). Additionally, we did not find a significant association between autophagosome degradation (SQSTM1) levels and memory impairment in females and males (Figure S1C-F).

**Figure 5.**
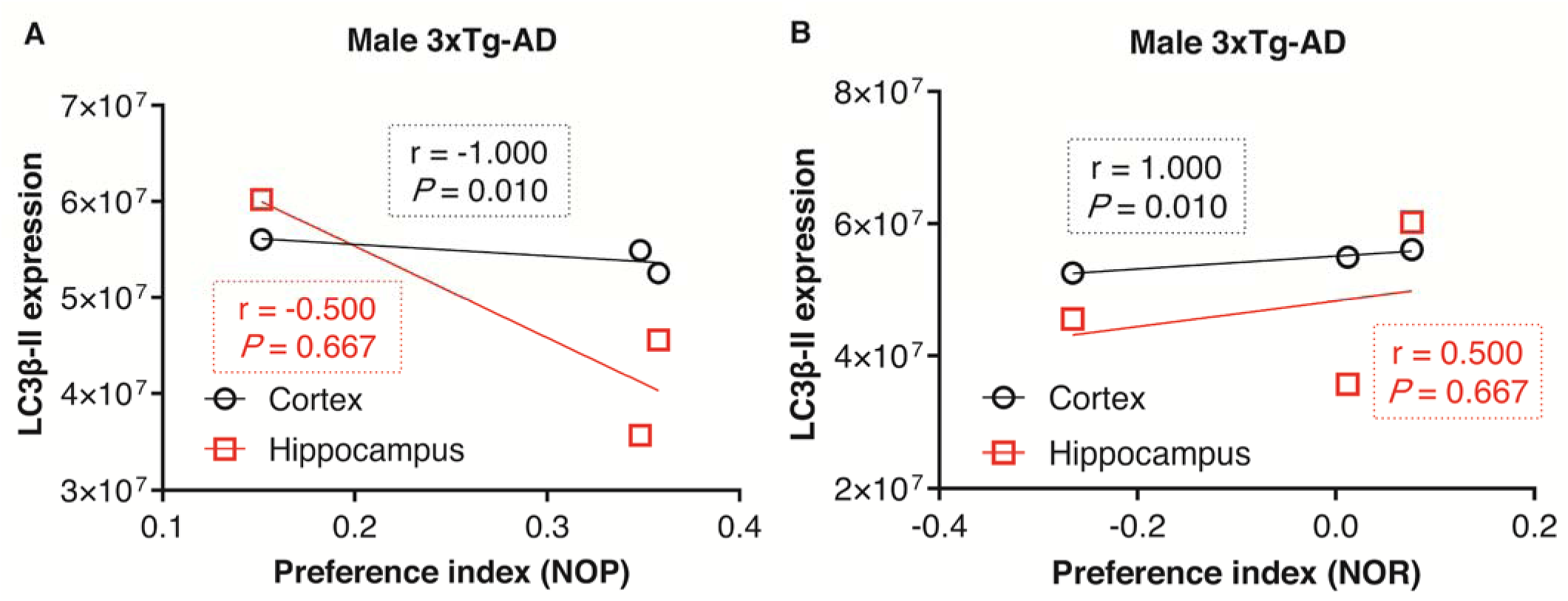
Association of Autophagosomes with Memory Impairment. **(A, B)** Association of LC3B-II levels (autophagosomes) with the NOP and NOR test in males, respectively. In females, no association was found between autophagosome and spatial memory impairment (P 0.05; see the supplement). In male mice, a negative correlation was observed between LC3B-II levels in the cortex and the NOP test, suggesting lower autophagosome levels are associated with better memory performance (NOP) (P = 0.010). Conversely, Higher LC3B-II levels in the cortex of male 3xTg-AD are associated with greater memory (P = 0.010). Spearman’s correlation test was used for statistical analysis, with P-value < 0.05 considered significant. LC3B-II, microtubule-associated protein 1 light chain 3 beta-II; NOR, novel object recognition; NOP, novel object placement.

We also performed correlation analyses between behavior findings and the BCL2-family proteins involved in mitophagy, including BNIP3L, BNIP3, and BCL2L13, as well as mitochondrial and mitophagosome numbers in the cortex and hippocampus of female and male 3xTg-AD mice. Our results showed that higher hippocampal BCL2L13 levels in females (r = 0.900, P = 0.037; Figure 6A) were correlated with greater non-spatial memory in the NOR test. Similarly, BNIP3L dimer levels in the cortex of females were positively correlated with spatial memory impairment (r = 0.900, P = 0.037; Figure 6B), while BNIP3L monomer expression levels were not associated with any behavior findings (Figure S2A-D).

**Figure 6.**
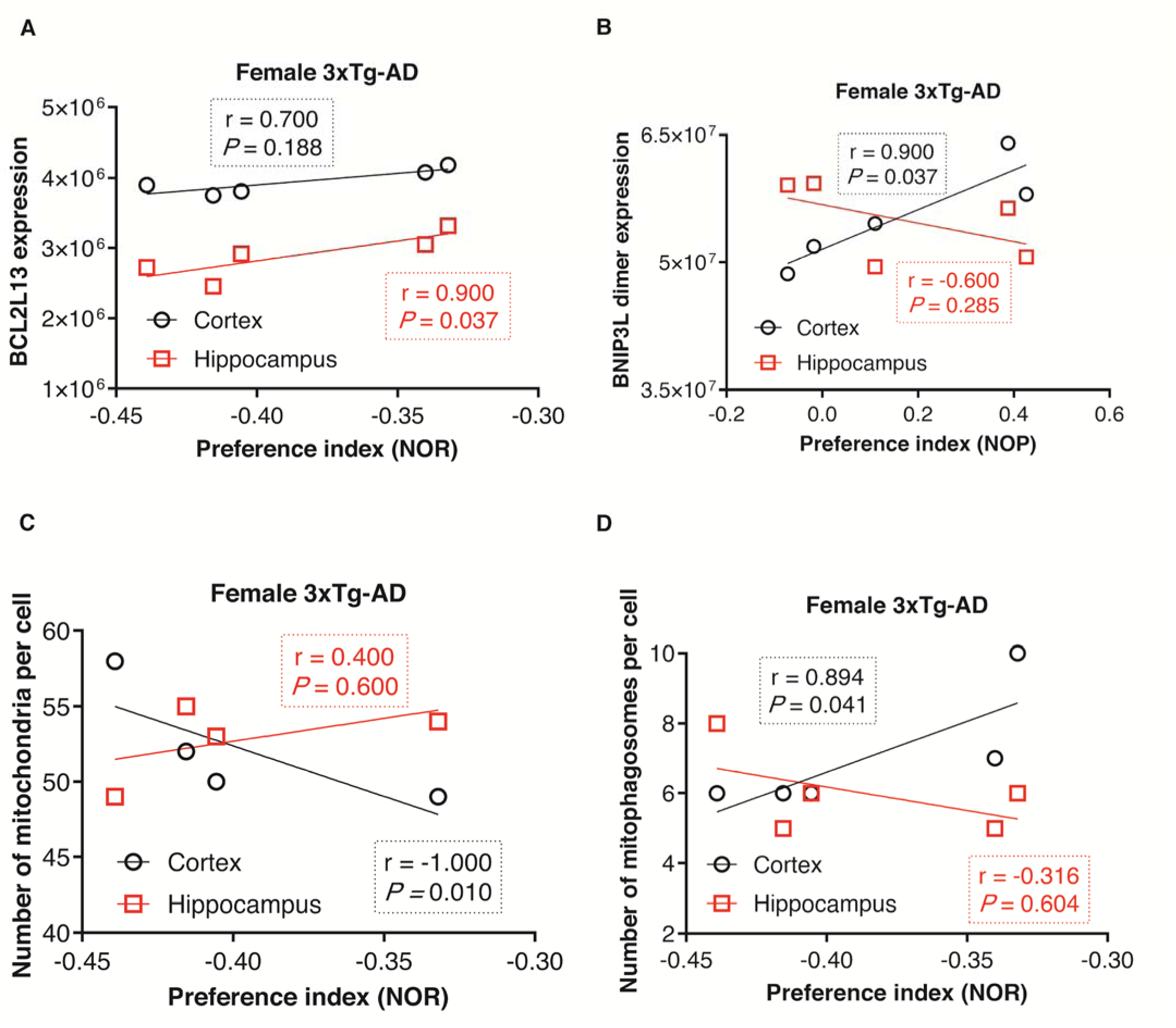
Association of BCL2 Family Proteins Expression Involved in Mitophagy and Mitochondria and Mitophagosome Numbers with Memory Impairment. **(A)** Correlation of NOR test with the levels of BCL2L13 in the cortex and hippocampus of females. Our results show that higher hippocampal expression of BCL2L13 is correlated with greater non-spatial memory in 3xTg-AD females (P = 0.037) **(B)** Correlation of NOP test with the levels of BNIP3L dimer in the cortex and hippocampus of females. In female 3xTg-AD mice, the NOP test showed a significant positive correlation with BNIP3L dimer levels in the cortex (P = 0.037). **(C)** Correlation of NOR test with mitochondria number in the cortex and hippocampus of female 3xTg-AD mice. Analysis shows a lower mitochondria number in the cortex correlated with greater non-spatial memory in female 3xTg-AD mice (P = 0.010). (**D)** Correlation of NOR test with mitophagosome number in the cortex and hippocampus of female 3xTg-AD mice. The number of mitophagosomes in the cortex was significantly positively correlated with the NOR behavior of females (P = 0.041). Spearman’s correlation test was used for statistical analysis, with a P-value < 0.05 considered statistically significant. BNIP3L, BCL2/adenovirus E1B interacting protein 3-like; BCL2L13, BCL2 like 13; NOR, novel object recognition; NOP, novel object placement.

Our results also demonstrated that the NOR behavior of female 3xTg-AD mice was negatively correlated with the number of mitochondria in the cortex (r = -1.000, P = 0.010; Figure 6C), but positively correlated with mitophagosomes number (r = 0.894, P = 0.041; Figure 6D). This suggests a possible link between mitophagy in cortex neurons and cognitive decline in female 3xTg-AD mice. No significant correlations were observed between mitophagosome and mitochondrial numbers in the cortex and hippocampus of male subjects with memory (Figure S3A-B). Furthermore, the number of mitochondria and mitophagosomes in both the cortex and hippocampus were not significantly related to the NOP test in females and males (Figure S3 C-F).

### Prediction of Mitophagosomes and Mitochondria in Cortical and Hippocampal Regions

Leveraging the behavioral data, we applied **machine learning** to predict the number of **mitophagosomes** and **mitochondria** in both the cortex and hippocampus, aiming to bridge cognitive behavior with underlying cellular dynamics. The prediction framework demonstrated favorable prediction accuracy in the number of mitophagosomes in the cortex and hippocampus and the number of mitochondria in the cortex (Figure 7A-C). However, the prediction for the number of hippocampus mitochondria based on behavior data was not meaningful (Figure 7D). This could be due to the limited sample size and the lack of a relationship between the number of hippocampal mitochondria and behavioral data (P > 0.32, Spearman correlation test).

**Figure 7.**
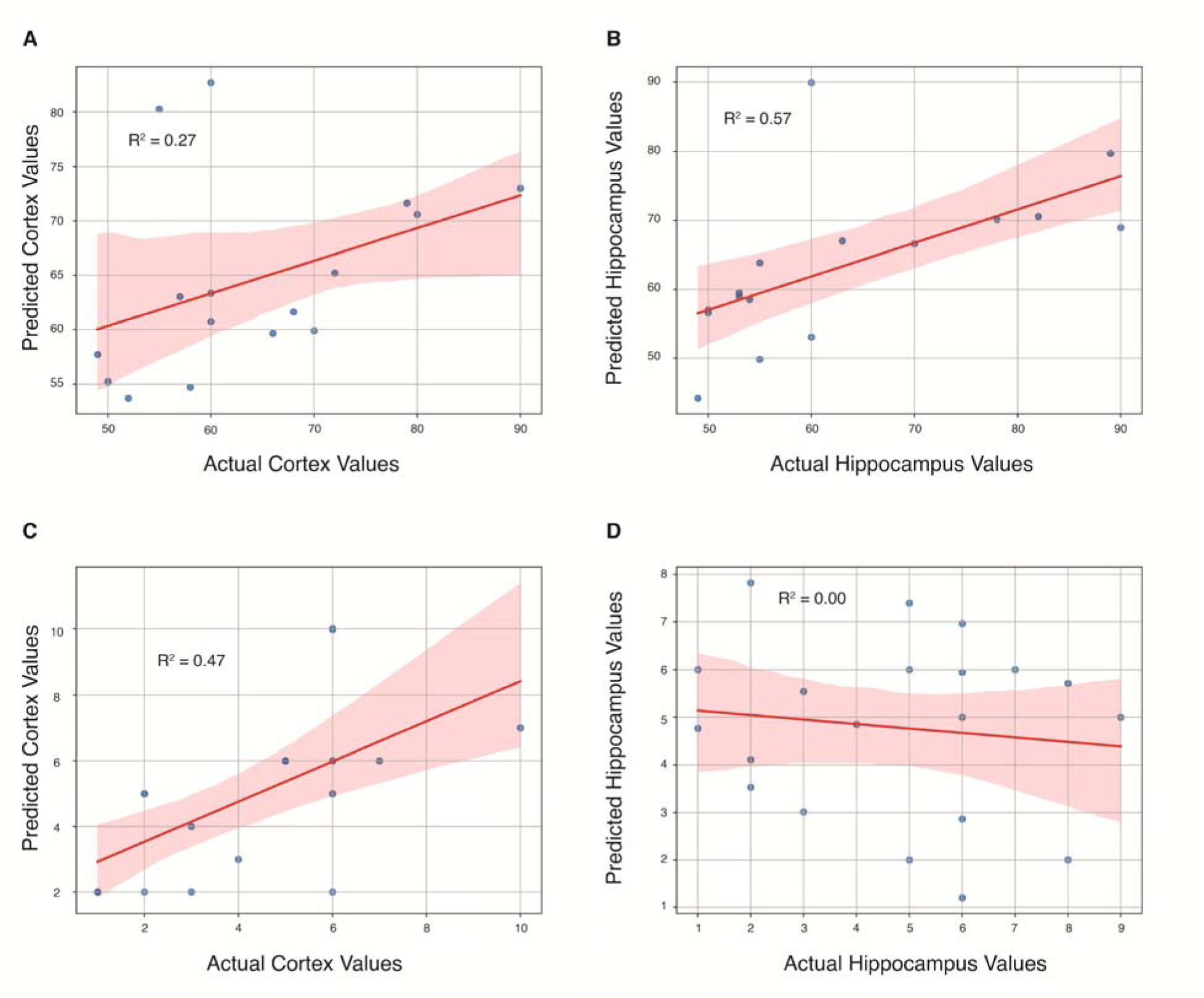
Scatter plots of predicted Cortex and Hippocampus values are shown as a function of their respective actual values, based on behavioral features coupled with machine learning. **(A, B)** Prediction of mitophagosomes in the cortex and hippocampus. **(C, D)** Prediction of mitochondria in the cortex and hippocampus. Favorable prediction accuracy for the number of mitophagosomes in the cortex and hippocampus, as well as for the number of mitochondria in the cortex.

## Discussion

Our study provides essential insights into the roles of autophagy and mitophagy in AD, with a specific focus on sex-specific differences that may influence the progression of the disease. Dysregulation of autophagy and mitophagy has been increasingly recognized as a significant contributor to AD pathogenesis, particularly through the accumulation of amyloid-beta plaques and MAPT/tau tangles, hallmark features of the disease. (43–49).

In this investigation, we identified sex-specific variations in key autophagy markers, particularly LC3B-II and SQSTM1, in the cortex and hippocampus, two brain regions critically involved in cognitive functions (50–52). Specifically, in female 3xTg-AD mice, LC3B-II levels were significantly elevated in the cortex, indicating autophagosome accumulation due to impaired autophagosome degradation, as supported by elevated SQSTM1 levels. In contrast, male 3xTg-AD mice exhibited reduced LC3B-II levels with no significant change in SQSTM1 in the hippocampus and cortex, suggesting fewer autophagosomes, but no major change in degradation pathways. This finding suggests that autophagic flux—the efficiency of autophagy—may be differentially regulated between males and females, potentially contributing to the observed differences in AD progression (53). The differential regulation of these markers may also reflect a more pronounced attempt at a compensatory autophagic response in females due to greater mitochondrial or protein damage, which may also contribute to sex differences in AD progression.

Mitophagy is critical in neurons, where mitochondrial dysfunction has been strongly linked to AD (54, 55). Mitochondria are essential for energy production and regulation of cellular metabolism, and their dysfunction can lead to increased oxidative stress and neuronal death, both of which are implicated in the neurodegenerative processes observed in AD (56, 57). Our study reveals significant sex-specific differences in mitophagy-related proteins, such as BNIP3L, BNIP3, and BCL2L13, across the cortex and hippocampus. For example, the higher levels of BNIP3L dimers observed in female 3xTg-AD mice suggest enhanced mitophagy activity, which might indicate a more robust response to mitochondrial damage (58, 59). This enhanced mitophagy activity is further supported by our observation of reduced mitochondrial numbers in the TEM analysis, indicating degradation defect in the female brain.

Conversely, the reduced expression of BNIP3 monomer in females with AD and the increased level of BNIP3 dimer in males suggest a potential impairment in mitophagy in females, leading to the accumulation of damaged mitochondria. This impairment could exacerbate oxidative stress and promote neurodegeneration, contributing to the observed sex differences in AD progression (60, 61). The differential regulation of BCL2L13, a protein involved in mitophagy, further highlights the complexity of these sex-specific differences. We observed higher BCL2L13 levels in the AD females, which was not seen in males. These patterns suggest that BCL2L13-dependent mitophagy pathways are differentially regulated between sexes, potentially contributing to the observed differences in mitochondrial turnover and AD progression.

In this study, we utilized two distinct behavioral tests to assess memory: NOR, which evaluates non-spatial memory, and NOP, which assesses spatial memory. These tests capture different facets of memory function—NOR focusing on recognition of object novelty, while NOP reflects memory for object location in space. This distinction provides a broader understanding of memory function by examining both spatial and non-spatial memory domains, offering insights into specific cognitive processes impacted by autophagy and mitophagy. Therefore, these molecular findings are corroborated by the behavioral outcomes observed in our study. The significant cognitive deficits displayed by female 3xTg-AD mice in the NOR test were closely associated with altered autophagy and mitophagy markers, particularly in the cortex. In females, the lack of significant associations between autophagy markers (LC3B-II, SQSTM1) and behavior suggests that mitochondrial health may play a more dominant role in determining cognitive outcomes, with a negative correlation observed between NOR performance and mitochondrial number in the cortex, and a positive correlation with mitophagosome number, indicating impaired mitochondrial turnover as a possible contributor to cognitive decline in AD. This highlights the importance of maintaining mitochondrial health in preventing or slowing the progression of neurodegenerative diseases (62, 63).

In contrast, in male mice, a positive correlation was observed between LC3B-II levels and NOR performance, suggesting that preserving autophagic flux could protect against cognitive decline. Notably, LC3B-II levels in males negatively correlated with NOP performance, hinting at a differential effect of autophagy on non-spatial and spatial memory. Additionally, higher hippocampal BCL2L13 levels in females were correlated with better NOR performance, while BNIP3L dimers in the cortex correlated with impaired NOP performance, potentially linking these mitophagy proteins to distinct aspects of memory in a sex-specific manner.

Overall, the observed differences between males and females in the correlations between autophagy and mitophagy markers with cognitive outcomes underscore the complexity of these mechanisms and suggest that mitochondrial quality and autophagic processes may impact memory domains in a sex-dependent manner

Autophagy and mitophagy play crucial roles in maintaining neuronal health, particularly in the cortex and hippocampus (64), which are key regions involved in cognition and memory. Dysregulation of these processes can lead to the accumulation of damaged proteins and organelles, contributing to neuronal dysfunction and cognitive decline (49). Studies have shown that impaired autophagy and mitophagy are associated with the accumulation of amyloid-beta and MAPT/tau pathology, which are linked to synaptic dysfunction and memory deficits in Alzheimer disease (65, 66). Specifically, in the hippocampus, a region critical for learning and memory, autophagy failure has been correlated with impaired synaptic plasticity, leading to deficits in spatial memory and learning (67, 68). In the cortex, autophagy and mitophagy dysfunction can affect the processing and integration of cognitive information, contributing to the behavioral and cognitive symptoms observed in Alzheimer disease (64). This underscores the importance of these cellular processes in the pathophysiology of AD and their potential as therapeutic targets.

Our study also demonstrates the potential of machine learning as a powerful tool for predicting disease progression and tailoring therapeutic interventions in AD. By integrating complex molecular and behavioral datasets, we achieved high prediction accuracy for mitophagosome and mitochondrial numbers in the cortex and hippocampus. This approach not only enhances our understanding of the relationships between autophagy, mitophagy, and cognitive function but also provides a framework for identifying potential biomarkers of AD progression.

The application of machine learning in this context is particularly promising for the development of personalized medicine in AD. Predictive models could help identify patients most likely to benefit from therapies targeting autophagy and mitophagy pathways, leading to more effective and individualized treatment strategies. As our understanding of the molecular underpinnings of AD continues to grow, the integration of machine learning and other advanced computational tools will be crucial in translating these findings into clinical practice (69, 70).

In conclusion, our study highlights the significant sex-specific differences in autophagy and mitophagy pathways in AD, with females showing increased autophagosome accumulation and enhanced mitophagy, particularly in the cortex, while males exhibit fewer autophagosomes and increased BNIP3-mediated mitophagy. These findings suggest that targeting autophagy and mitophagy pathways in a sex-specific manner could lead to more effective therapeutic interventions for AD. The differential regulation of these processes between males and females underscores the need for personalized therapeutic approaches that consider the unique molecular and behavioral profiles of each patient. Additionally, the integration of machine learning into AD research offers a promising avenue for developing predictive models that could guide future therapeutic interventions. As we continue to explore the intricate connections between autophagy, mitophagy, and AD, our findings pave the way for new strategies aimed at mitigating the impact of this devastating disease.

## Contribution

**Aida Adlimoghaddam** performed the experiments, prepared the initial draft of the paper, and reviewed the final manuscript. **Fariba Fayazbakhsh**, **Mohsen Mohammadi**, **Amir Barzegar Behrooz**, **Teng Guan**, and **Farhad Tabasi** contributed to the preparation of the manuscript draft and finalized the figures. **Mahmoud Aghaei** and **Zeinab Babaei** conducted all correlation analyses. **Iman Beheshti** performed the machine learning analysis. **Daniel J. Klionsky** reviewed and finalized the manuscript, providing significant insights on autophagy and mitophagy mechanisms. **Benedict Albensi** contributed to the finalization of the manuscript and provided expertise on Alzheimer’s disease. **Saeid Ghavami** supervised, led, and designed the project and was responsible for the final review and approval of the manuscript.

## Abbreviations

Aβ: amyloid-β
AD: Alzheimer disease
NOP: novel object placement
NOR: novel object recognition
TEM: transmission electron microscopy

