## Supplementary Figures for "Sex and Region-Specific Disruption of Autophagy and Mitophagy in Alzheimer’s Disease: Linking Cellular Dysfunction to Cognitive Decline"

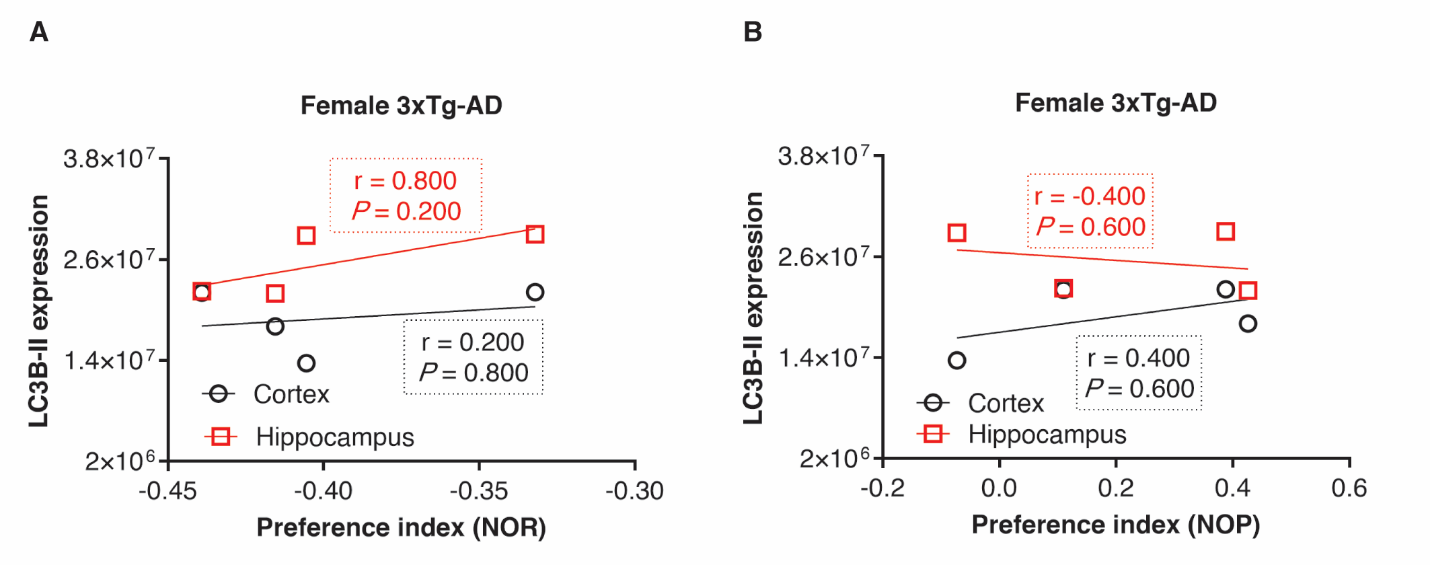

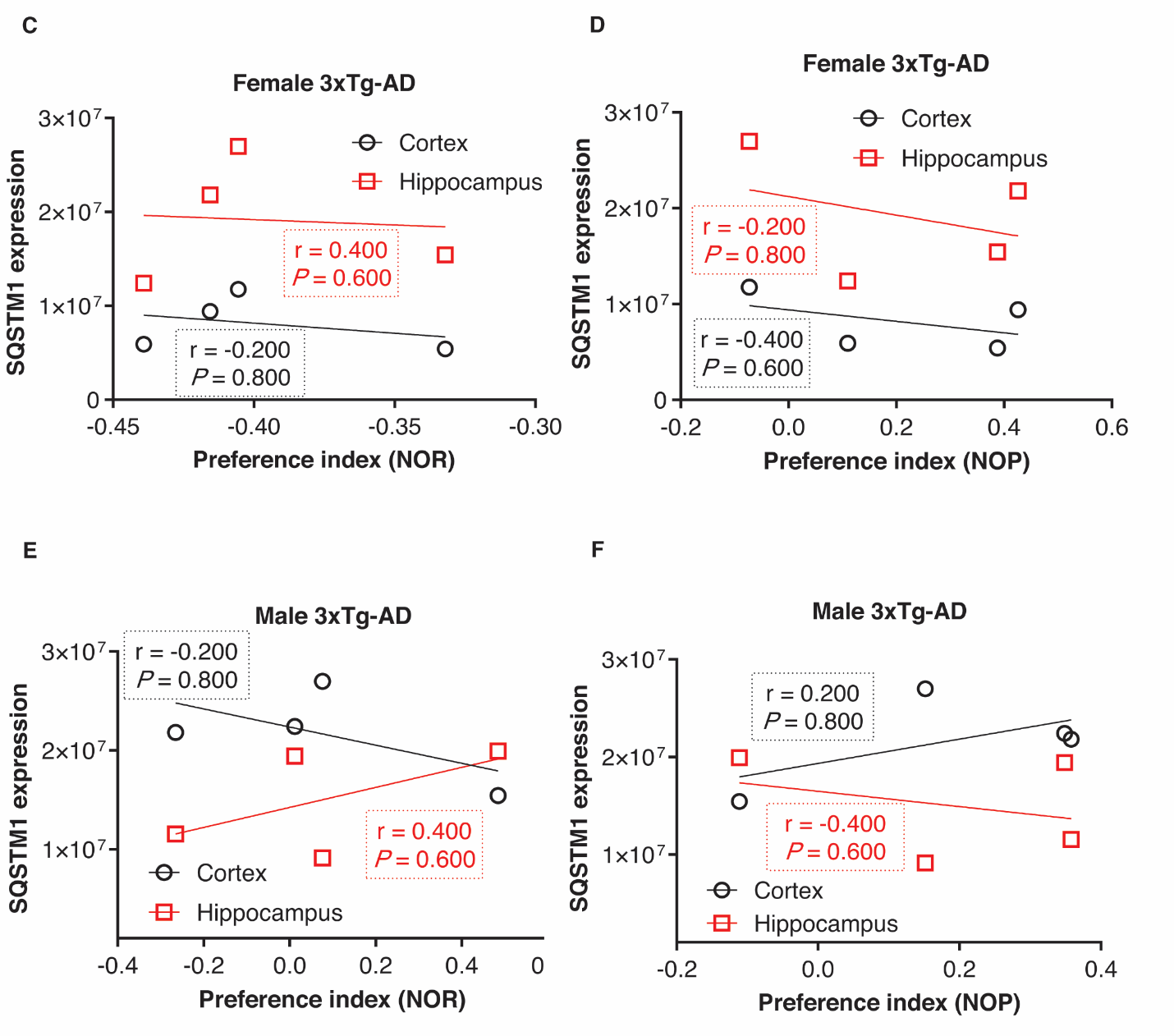

**Figure S1. Association of Autophagosome with Memory Impairment** **(A, B)** Association of LC3B-II levels (autophagosome) with the NOR and NOP test in females, respectively. In females, no association was found between autophagosome and memory deficit (*P* ˃ 0.05). **(C, D)** Autophagosome degradation (SQSTM1) level association with the NOP test in females and males. Our analyses reveal no significant association between autophagosome degradation levels and the NOP test in 3xTg-AD mice (*P* ˃ 0.05). **(E, F)** SQSTM1 level association with the NOR test in females and males. Our analyses reveal no significant association between SQSTM1 levels and NOR test in 3xTg-AD mice (*P* ˃ 0.05). Statistical analyses were conducted using Spearman's correlation test, with a *P*-value < 0.05 considered statistically significant. LC3B-II, microtubule-associated protein 1 light chain 3 beta-II; SQSTM1, sequestosome 1; NOR, novel object recognition; NOP, novel object placement.

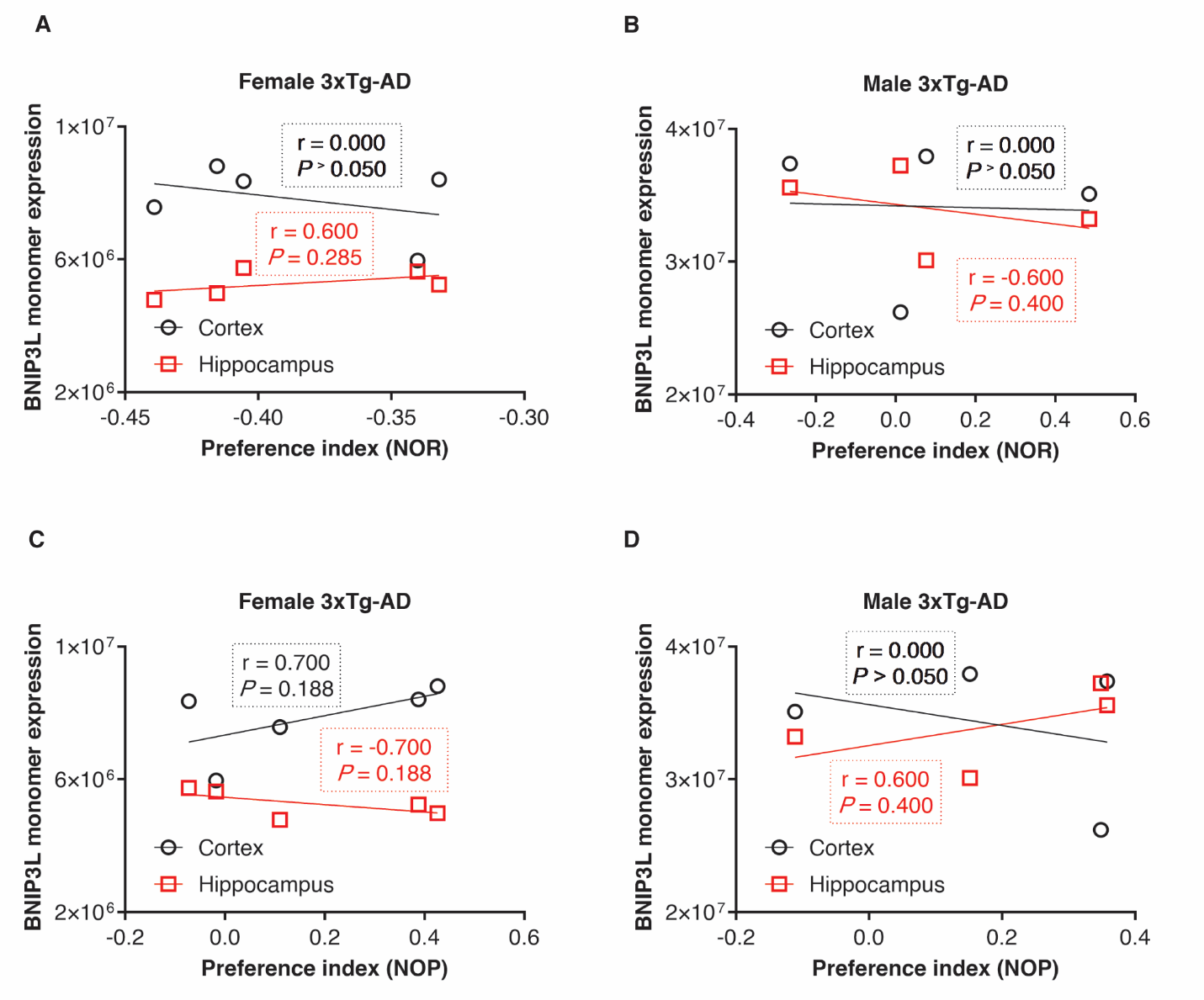

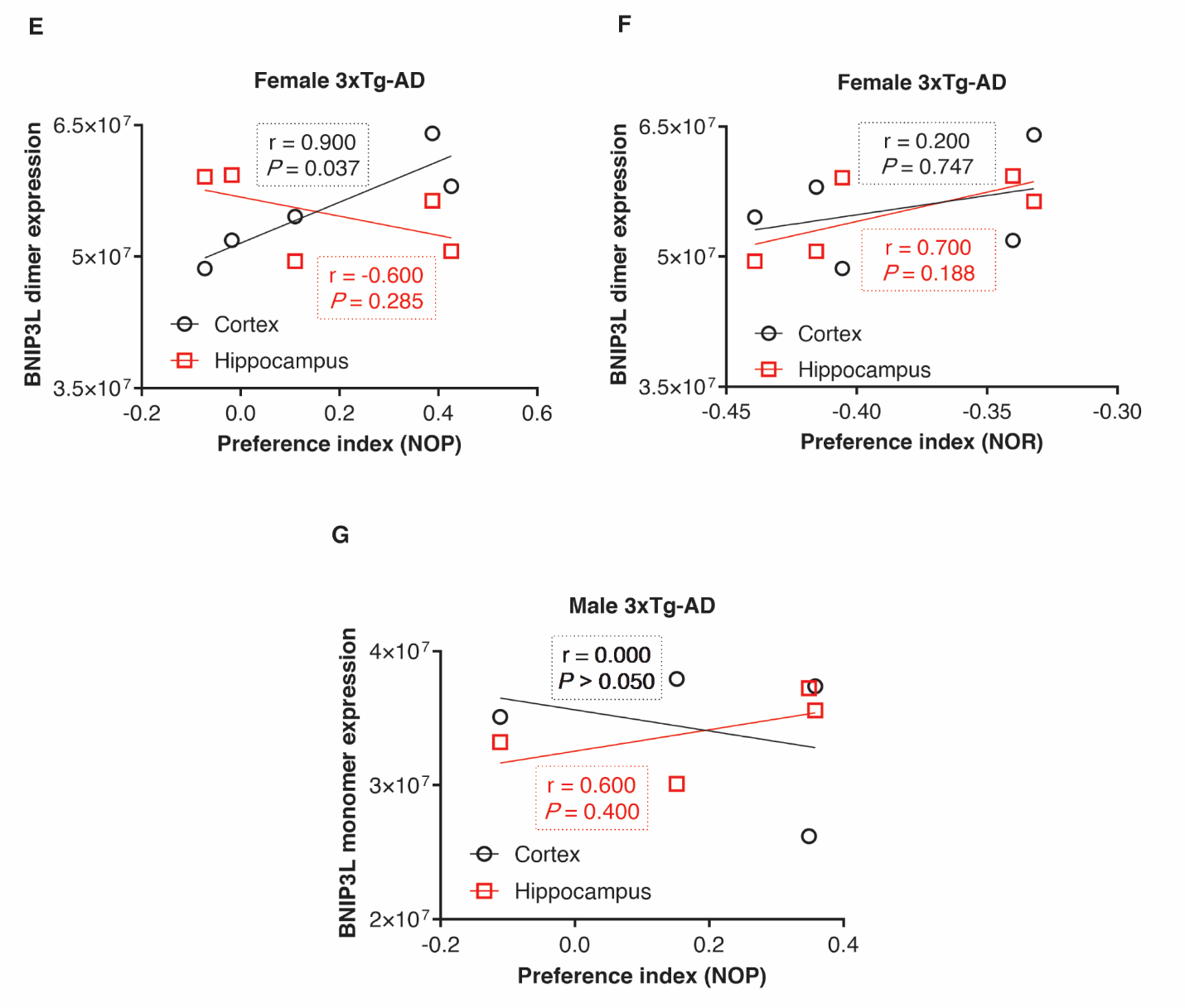

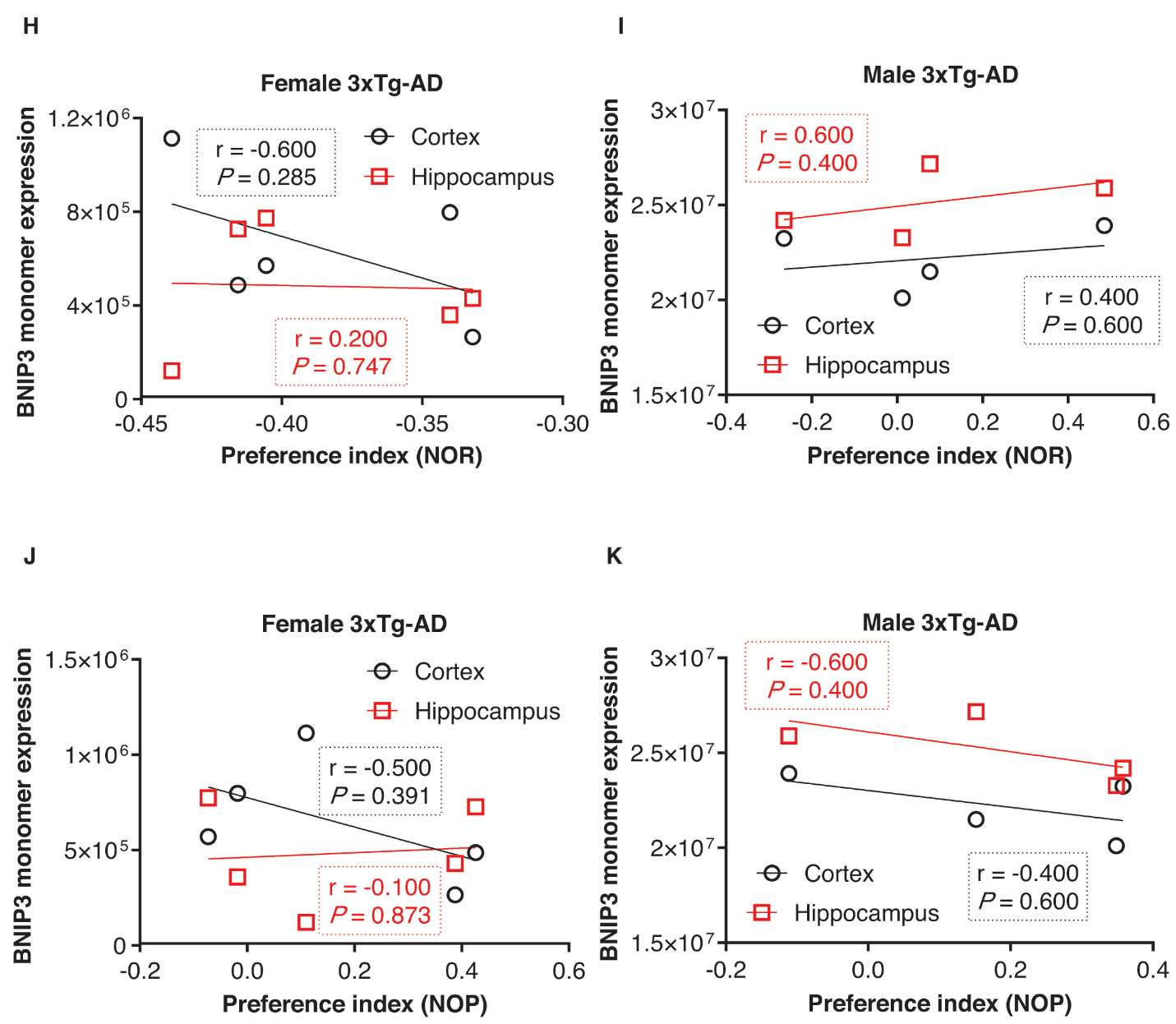

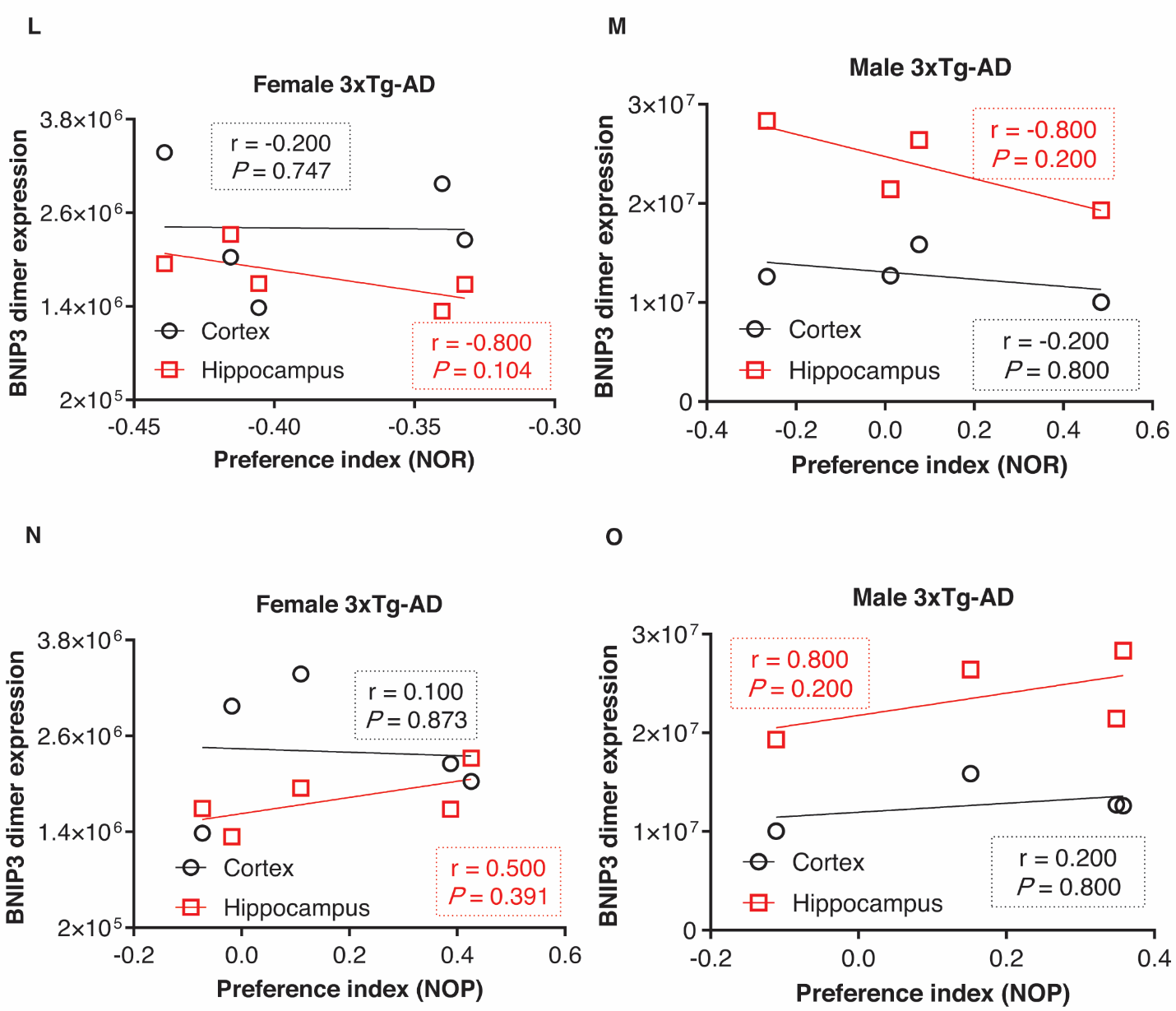

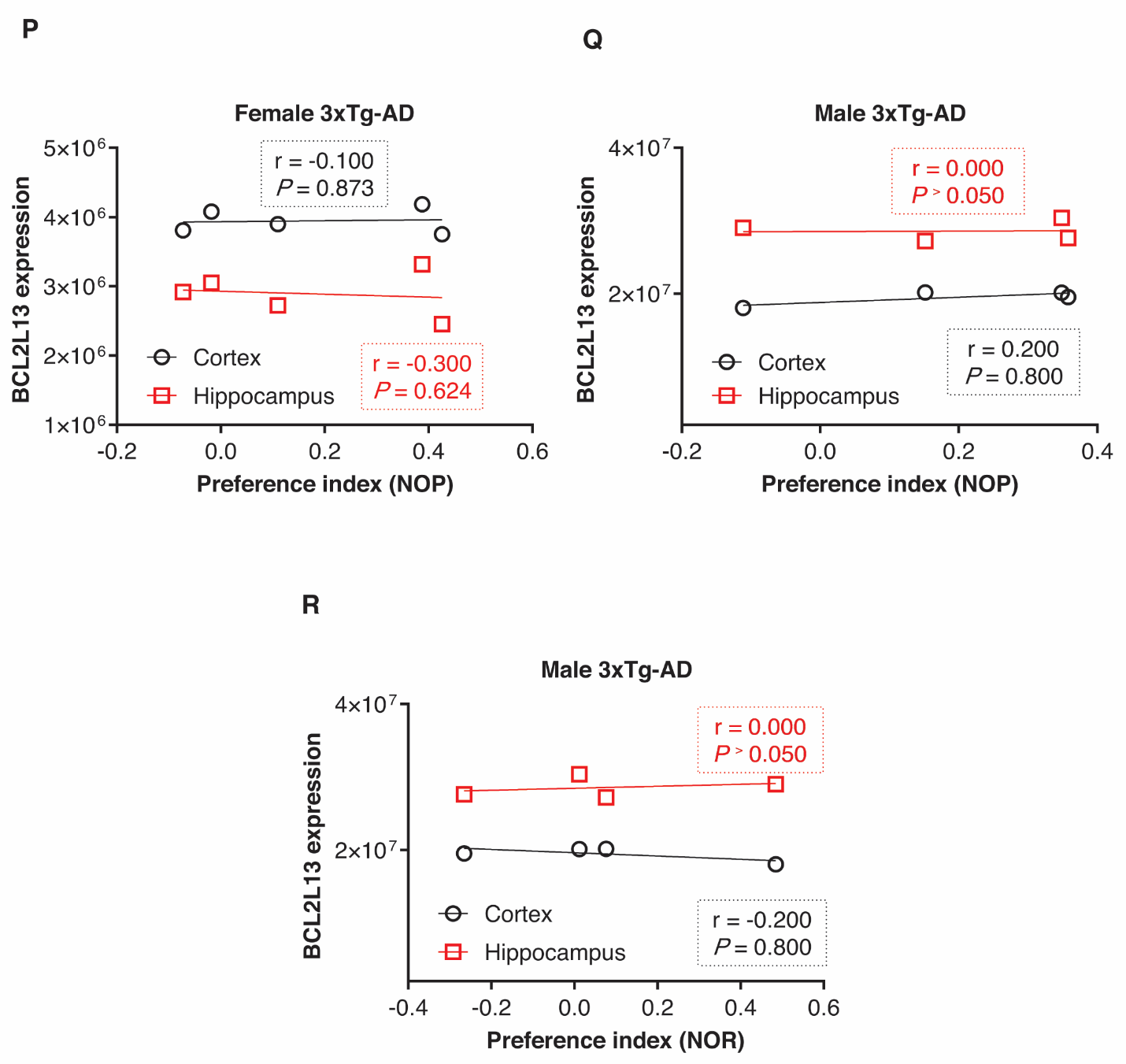

**Figure S2. Association of BCL2-Family Proteins Expression Involved in Mitophagy with Memory Impairment** **(A, B)** Correlation of NOR test with the levels of BNIP3L monomer in the cortex and hippocampus of females and males. **(C, D)** Correlation of NOP test with the levels of BNIP3L monomer in the cortex and hippocampus of females and males. **No association was found between the monomer forms of** BNIP3L **and memory impairment (*P* ˃ 0.05). (E, F)** Correlation of BNIP3L dimer with NOR test in the cortex and hippocampus of females and males. **No association was found between the dimer forms of** BNIP3L **and cognition memory deficit (*P* ˃ 0.05).**  **(G)** Correlation of BNIP3L dimer with NOP test in the cortex and hippocampus of males. Analysis shows no association of spatial memory with BNIP3L expression (*P* ˃ 0.05). **(H, I)** Correlation of NOR test with the expression levels of BNIP3 monomer in the cortex and hippocampus of female and male 3xTg-AD mice. **(J, K)** Correlation of NOR test with the expression levels of BNIP3 dimer in the cortex and hippocampus of male and female 3xTg-AD mice. **(L, M)** Correlation of NOP tests with the expression levels of BNIP3 monomer in the cortex and hippocampus of female and male 3xTg-AD mice. **(N, O)** Correlation of NOP test with the expression levels of BNIP3 dimer in the cortex and hippocampus of female and male 3xTg-AD mice. Our investigation shows no significant correlation between monomer and dimer of BNIP3 expression (mitophagy) and memory deficit (*P* ˃ 0.05). **(P, Q)** Correlation of BCL2L13 levels with NOP test in females and males. Our analysis shows no association between BCL2L13 levels and memory deficit (*P* ˃ 0.05). **(R, S)** Correlation of BCL2L13 levels with NOR test in male. No correlation was found between BCL2L13 and cognition memory in males (*P* ˃ 0.05). Statistical analyses were conducted using Spearman's correlation test to determine statistical significance, with a *P*-value < 0.05 considered statistically significant. BNIP3L, BCL2/adenovirus E1B interacting protein 3-like; BNIP3, BCL2/adenovirus E1B interacting protein 3; BCL2L13, BCL2 like 13; NOR, novel object recognition; NOP, novel object placement.

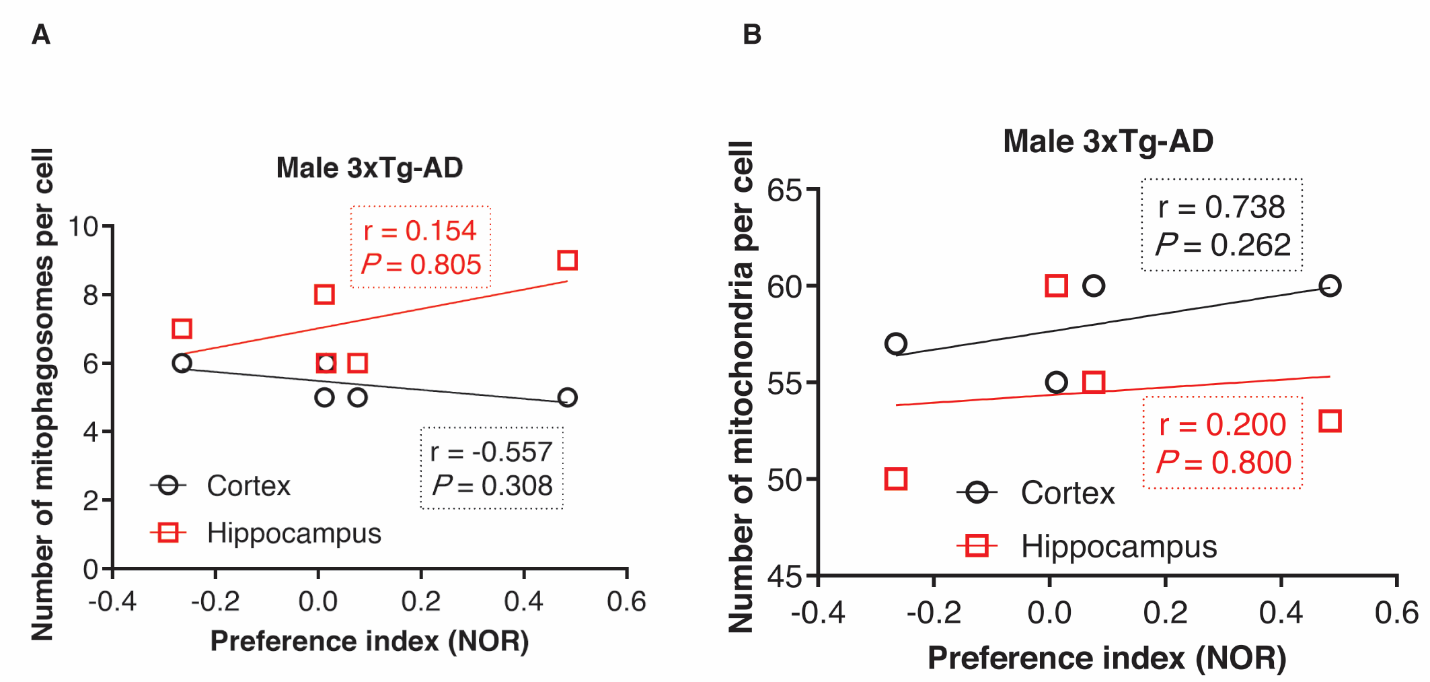

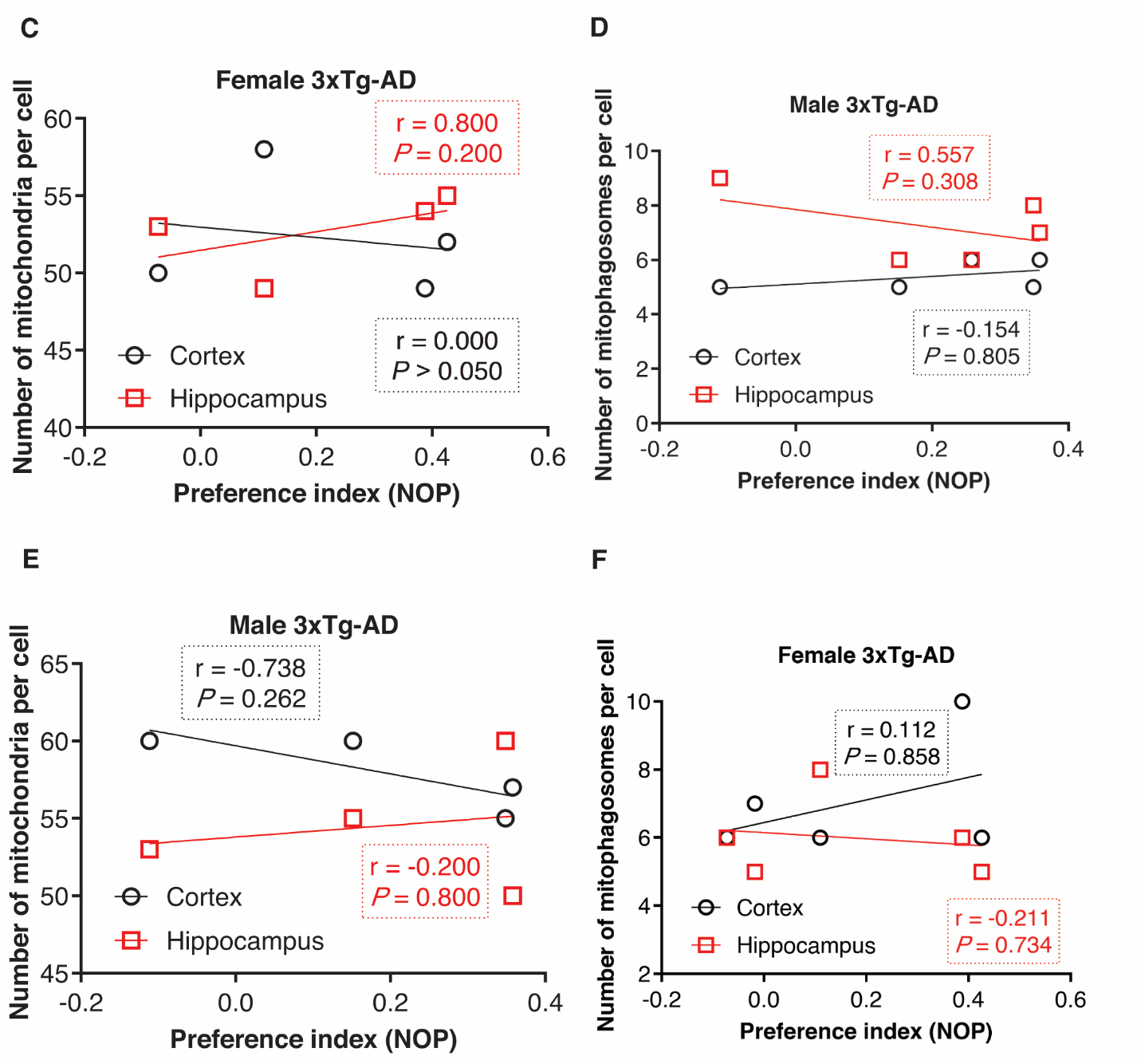

**Figure S3. Association of Mitochondria and Mitophagosome Numbers with Memory Impairment**. **(A, B)** Correlation of NOR test with mitophagosome and mitochondria number in the cortex and hippocampus of male 3xTg-AD mice. Results show no correlation between mitophagosome and mitochondria number and cognition memory in male mice (*P* ˃ 0.05). **(C, D)** Correlation of NOP test with mitophagosome number in the cortex and hippocampus of female and male 3xTg-AD mice. **(E, F)** Correlation of NOP test with mitochondria number in the cortex and hippocampus of female and male 3xTg-AD mice. Our investigation showed no significant association of mitochondria and mitophagosome number with spatial memory (*P* ˃ 0.05). Statistical significance was tested using Spearman’s correlation test. A *P*-value < 0.05 was considered statistically significant. NOR, novel object recognition; NOP, novel object placement.
